# Developmental landscape of human forebrain at a single-cell level unveils early waves of oligodendrogenesis

**DOI:** 10.1101/2021.07.22.453317

**Authors:** David van Bruggen, Fabio Pohl, Christoffer Mattsson Langseth, Petra Kukanja, Hower Lee, Mukund Kabbe, Mandy Meijer, Markus M. Hilscher, Mats Nilsson, Erik Sundström, Gonçalo Castelo-Branco

## Abstract

Oligodendrogenesis in the human central nervous system has been mainly observed at the second trimester of gestation, a much later developmental stage compared to mouse. Here we characterize the transcriptomic neural diversity in the human forebrain at post conceptual weeks (PCW) 8 to 10, using single-cell RNA-Seq. We find evidence of the emergence of a first wave of oligodendrocyte lineage cells as early as PCW 8, which we also confirm at the epigenomic level with single-cell ATAC-Seq. Using regulatory network inference, we predict key transcriptional events leading to the specification of oligodendrocyte precursor cells (OPCs). Moreover, by profiling the spatial expression of fifty key genes using In Situ Sequencing (ISS), we identify regions in the human ventral fetal forebrain where oligodendrogenesis first occurs. Our results indicate evolutionary conservation of the first wave of oligodendrogenesis between mouse and human and describe regulatory mechanisms required for human OPC specification.

## Introduction

The understanding of diversification of cell types has been a long outstanding challenge in biology. The human brain is one of the most diverse tissues in the mammalian body, and on the evolutionary scale, the emergence of the first neuronal cell-types dates back millions of years (Arendt et al., 2016a). Recent advances in single-cell transcriptomics allow capture of this diversity of cell states in developing tissues (Buenrostro et al., 2015; Cao et al., 2019; Hochgerner et al., 2017; Islam et al., 2014; Klein et al., 2015; La Manno, 2019; Marioni and Arendt, 2017). One of the most defining characteristics of primate evolution, and specifically human evolution is cortical expansion (Polioudakis et al., 2019; de la Torre-Ubieta et al., 2018; Won et al., 2016; Zhu et al., 2018). The human cortex is much larger and possibly more complex compared to any animal. However, the mammalian brain contains the most diverse collection of known cell-types compared to any other tissue. And as such, it has been studied intensely. From an evolutionary perspective, many aspects of human brain development are similar to other mammalian and even other vertebrates (Arendt et al., 2016b; Houart et al., 1998, 1998; La Manno et al., 2016). Transcription factor families, such as the basic helix-loop-helix (bHLH) and forkhead-box (FOX) amongst others have evolved and diversified forming a web of interweaved dependencies in the spatio-temporal context of brain development. Dorsal-ventral patterning factors, morphogens, chromatin modifying enzymes, and regulatory elements play a role in the development of the brain across species. Moreover, during the evolution of the vertebrate and then mammalian central nervous system (CNS) glial cells appeared. Glial cells such as astrocytes and oligodendrocytes have been traditionally thought to have mainly supportive roles to neurons in the CNS. However, recent findings indicate that glial cells are involved in many other functions in the CNS (Di Bella et al., 2021; Falcão et al., 2018; Kirby et al., 2019).

The formation of glial cells during mouse development occurs after neurogenesis (Rowitch and Kriegstein, 2010). Three main waves of oligodendrogenesis have been characterized in the mouse brain, with the first starting at embryonic day (E)12.5 from ventral regions (Kessaris et al., 2006). We have recently shown that oligodendrogenesis in mouse entails an intermediate stage of pre-OPCs, between neural progenitors and OPCs (Marques et al., 2018). During human fetal development, OPCs have been mainly observed at the second trimester of gestation, with the first cells detected at 16 weeks, and much larger numbers detected at 22 weeks (Huang et al., 2020; McClain et al., 2012; Sim et al., 2011; Windrem et al., 2004). These OPCs can be isolated with antibodies targeting CD140a/PDGFRA, can originate both oligodendrocytes and astrocytes and are highly migratory and myelinogenic upon transplantation to the mouse brain (Sim et al., 2011; Windrem et al., 2020). More recently, pre-OPCs (Marques et al., 2018) were also observed in the human cortex at 20-24 post conception weeks (PCW) (Huang et al., 2020). These second trimester pre-OPCs might correspond to the last cortical wave as described in mouse (Di Bella et al., 2021; Kessaris et al., 2006; Winkler et al., 2018). They exhibit as hallmark the expression of EGFR (Huang et al., 2020). Nevertheless, it is unclear whether earlier distinct waves of oligodendrogenesis occur in human. Interestingly, OLIG1+/PDGFRA+ cells might arise in the forebrain at as early as 9-10 weeks of gestation (Jakovcevski and Zecevic, 2005). Here we used single cell transcriptomics, epigenomics (UCSC Cell Browsers (Speir et al., 2021) for visualization available at https://human-forebraindev.cells.ucsc.edu) and spatial transcriptomics to characterize the specification of glial lineages during early human forebrain development, at PCW 8-11. We found that as early as PCW 8 radial glial cells start being specified in the ventral forebrain into the oligodendrocyte lineage, indicating that a first wave of oligodendrogenesis indeed occurs early in the first trimester of human development.

## Results

### Single-cell RNA-seq of human fetal forebrain at the first trimester uncovers early cell transitions from neural progenitors towards neuronal fates, but also glial fates

We performed droplet-based single-cell RNA-seq on human fetal forebrain tissue at the first trimester, on 5 separate samples from 4 different fetuses of PCW 8, 8.5, 9, and 10, using the 10x Genomics platform (versions 2 and 3). We obtained an average number of 7400 UMIs and 2500 expressed genes (Figure S1A). After quality control, we retained 25161 cells. We batch corrected the lower dimensional space using Harmony (Korsunsky et al., 2019) to integrate the v2 and v3 data, while retaining biological differences. We then clustered the data using Louvain clustering on the obtained Jensen-Shannon distance (JSD) matrix from the integrated PCA resulting in 32 clusters (Figure 1A, B, S1B, see Methods for details). By assessing the expression of known neural markers, we were able to identify populations of radial glial cells, neuroblasts, excitatory and inhibitory neurons, but also cells of glial, endothelial and vascular and leptomeninges (VLMCs) lineages (Figure 1A,B). These populations were enriched in specific time-points, reflecting different stages of development (Figure 1C, D, Figure S1C).

**Figure 1.**
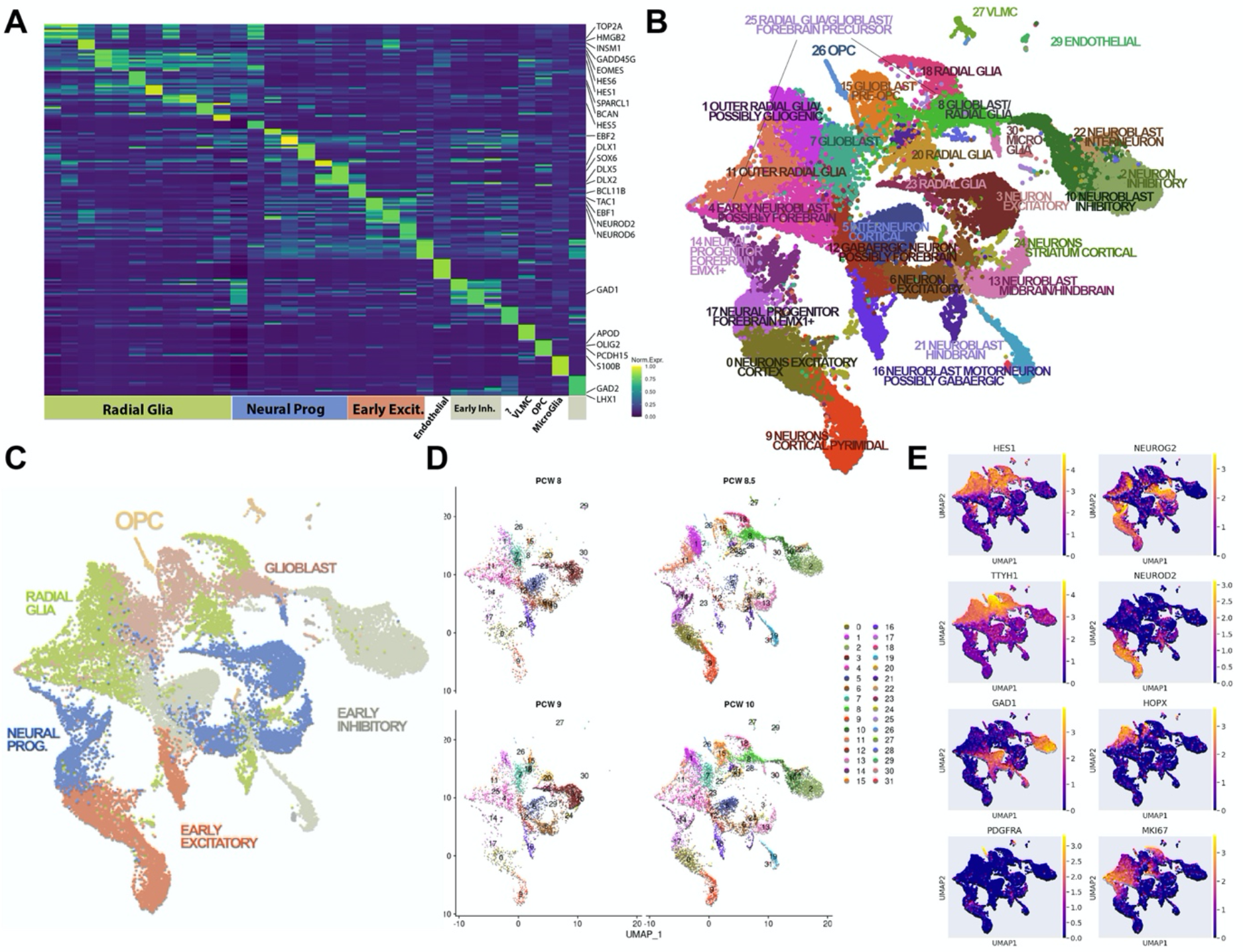
Single cell RNA-seq reveals a continuous manifold of developmental transitions into glial fates already at the human PCW 8-10 forebrain. A) Heatmap showing the top 10 genes enriched for each cluster. B) UMAP of 25161 cells depicting the clusters and their inferred identities based on marker expression. C) Coarse clusters showing the main developmental stages in the early brain. D) UMAP depicting the different post conception ages in the experiment. E) Representative genes for the coarse cluster, expression overlayed in UMAP from Panel B.

Clustering revealed populations of radial glial cells expressing genes such as Hes Family BHLH Transcription Factor 1 (*HES1*), Fatty Acid Binding Protein 7 (*FABP7*), Paired Box 6 (*PAX6*), and Vimentin (*VIM*) (Figure1A, E). The radial glial population could be further distinguished by the expression of outer radial glial genes such as HOP Homeobox (*HOPX*), a well-known marker for outer radial glia (Nowakowski et al., 2017; Pollen et al., 2015). Broadly we could resolve cycling cells expressing *MKI67*, and cells belonging to the early excitatory lineages, expressing genes such as Neuronal Differentiation 2 and 6 (*NEUROD2*, *NEUROD6*) among others. And early inhibitory lineages expressing genes such as Glutamate Decarboxylase 1 (*GAD1*), and *GAD2*, as well as LIM Homeobox 1 (*LHX1*) Figure 1A, E).

Moreover, we could also distinguish glioblast-like populations such as Cluster 15, expressing Brevican (*BCAN*), Regulatory Factor X4 (*RFX4*), ZFP36 Ring Finger Protein Like 1 (*ZFP36L1*), Cyclin D1 (*CCND1*), and MyoD Family Inhibitor (*MDFI*). We detected expression of the OL lineage transcription factor *OLIG2* and *NKX2-2* in this cluster (Figure S1D), which could indicate that early OL lineage cells might contribute to this cluster. Additionally, we found early astrocyte lineage cells in this cluster expressing Secreted Protein Acidic And Cysteine Rich (*SPARC*), TSC22 Domain Family Member 4 (*TSC22D4*), Endothelin Receptor Type B (*EDNRB*), SPARC Like 1 (*SPARCL1*), and Glial Fibrillary Acidic Protein (*GFAP*) (Figure S1D). In sum, cluster 15 could be a population that has the potential to become OPC and astrocytes.

### Specification of human OPCs occurs already at PCW 8

Importantly, at these early stages we could resolve a population of cells expressing OL lineage genes such as Platelet Derived Growth Factor Receptor Alpha (*PDGFRA*), SRY-Box Transcription Factor 10 (*SOX10*), NK2 Homeobox 2 (*NKX2-2*), Oligodendrocyte Transcription Factor 1/2 (*OLIG1*), (*OLIG2*) and many other OPC specific genes (Figure 1A, E. Figure 2G, and Figure S1D). The OPC population (Cluster 26) was present in all timepoints (PCW 8-10, Figure S1C) and seemed to be closely related to the glioblast population (Cluster 15), both by co-expression of glial lineage genes such as *BCAN*, *TTYH1*, *ZFP36L1*, *HES1*, as well as expression of OL lineage genes such as OLIG2, SOX10, and NKX2-2 (Figure S1D). We named this cluster pre-OPCs (Marques et al., 2018) and proceeded to identify differentially expressed genes between the OPC and putative pre-OPC population (Wilcoxon rank-sum test). OPCs expressed genes strongly associated with OPC identity such as *S100B, KCND2, OLIG1, PCDH15, SCRG1, LHFPL3, OPCML, APOD*, and *OLIG2*, among other genes (Figure S1E). Conversely, the pre-OPC population expressed genes that indicated a progenitor identity, expressing genes such as *VIM, FABP7, TTYH1, SPARCL1, HES1, RMST, CLU*, and *FOS* (Figure S1F). Additionally, we observed differentially expressed transcription factors between the OPC and the pre-OPC identity (Figure S1F). OPC identity strongly expressed factors known to be important in the specification and maintenance of OPCs such as *OLIG1, OLIG2, SOX6, NKX2-2,* and *SOX10* (Jakovcevski and Zecevic, 2005; Rowitch and Kriegstein, 2010; Windrem et al., 2004). Importantly, several other transcription factors and co-factors that have not yet/or have been weakly implicated in OL specification were identified, including *LUZP2, NCALD, NR0B1, ETV1, MITF*, and *TRAF4* (Figure S1F). Conversely pre-OPC identity seemed to be associated with factors known to be important for glial cell formation and genes associated with the neuro-glial switch, migration, and activation such as *HES1, GLIS3, FOS, NFIA, NFIB, HES4, TSC22D4, NFATC2, JUNB, HES5*, and *FOXJ1* among others (Figure S1F).

**Figure 2.**
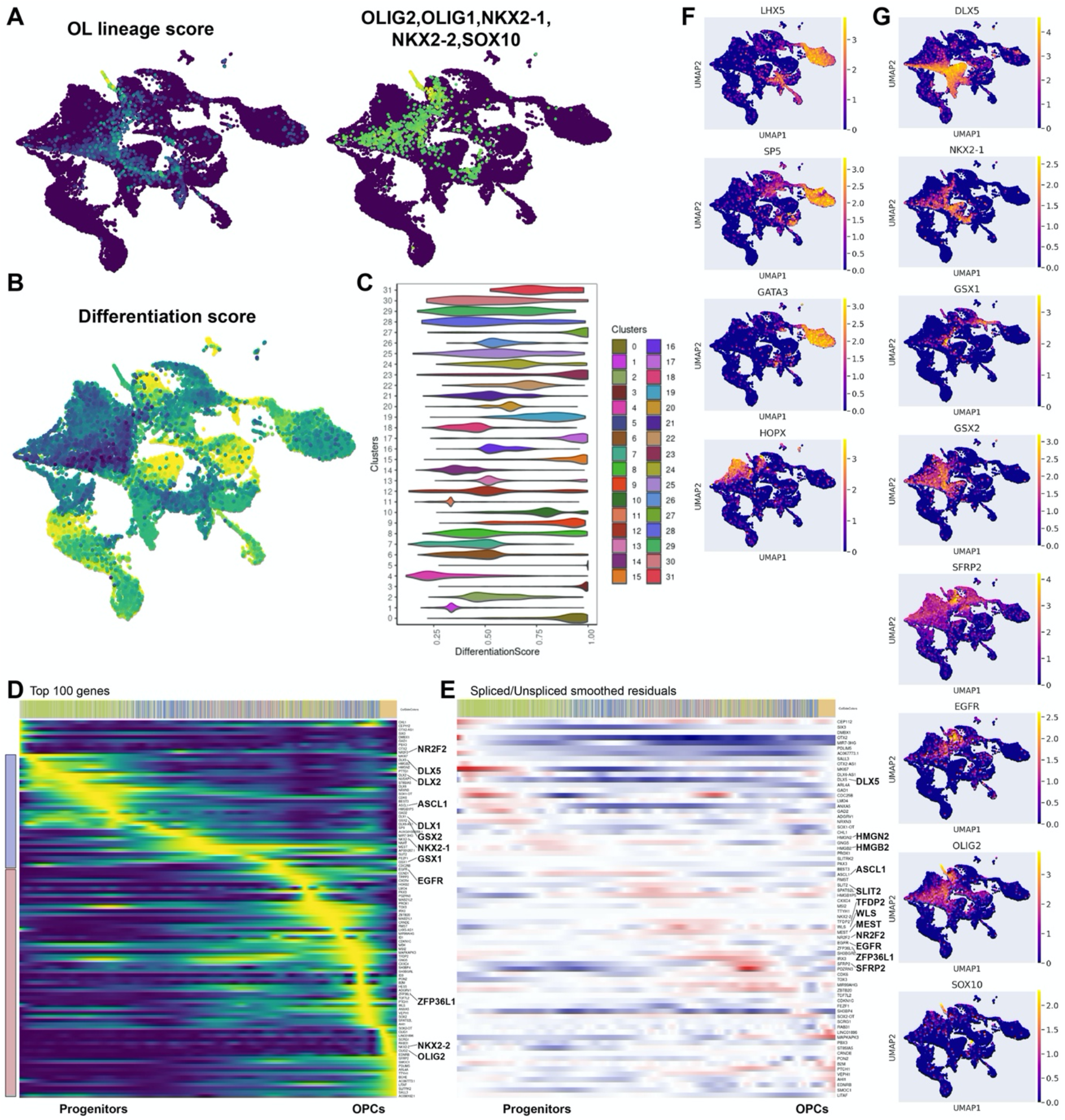
Convergent paths from radial glia, through glioblasts, to OPCs in the human PCW8-10 forebrain. A) UMAP showing the OPC lineage association score (Left), and the joint expression profile of the major OL lineage markers *OLIG2*, *OLIG1*, *NKX2-1*, *NKX2-2* and *SOX10* for comparison (Right) B) UMAP showing the estimated differentiation score of every cell in the dataset according to the lineage analysis over all lineages (Left). C) Violin plot showing estimated differentiation score for every cluster. Cluster number according to Figure 1B. D) Smoothed depiction of the top 100 genes correlating with the OL lineage, ordered over time. Color bar colored according to the coarse clustering. Blue color bar on the left highlights expression of early lineage genes mainly involving patterning genes. Red bar, genes expressed during and after the glioblast stage. E) The top 100 genes filtered for genes lacking both spliced and unspliced counts, showing the difference between the min/max normalized smoothed expression within the lineage cells, Red = positive residual (possible upregulation of gene), Blue = negative residual (possible downregulation of gene). Genes are ordered according to residual value. Color bar colored according to the coarse clustering. F) UMAPs depicting *LHX5*, *SP5*, *GATA3*, and *HOPX* expression, showing possible outer radial glia progenitor groups. G) UMAPs of patterning genes expressing along the OL lineage.

### Lineage inference by back-diffusion reveals two paths towards human OPCs, with outer radial glia and medial ganglionic eminence radial glia as likely origins

We next sought to confirm if we correctly inferred the pre-OPC state as the state OPCs transition through during OL fate consolidation in early human forebrain development. To avoid losing the gradients involved in differentiation, which might be broken by the Harmony algorithm, we reintegrated the V2 and V3 datasets using a customized approach inspired by the reciprocal PCA and anchor-based integration as found from Seurat version 3 on (Stuart et al., 2019) (see Methods). This necessitated re-clustering of the newly integrated data for lineage assessment leading to 30 clusters (Figure S2A). We then used the corrected PCA space as input for scVelo (Bergen et al., 2019) to obtain a transition matrix of cell velocities. We generated a diffusion map (Haghverdi et al., 2016) of the integrated PCA space to obtain a transition matrix of transcriptomic similarities, we performed canonical correlation analysis (CCA) on both these transition matrices as input to the final diffusion map to generate a manifold congruent with transcriptional and velocity distances that can capture non-linear relationships (see Methods).

We reasoned that the manifold of human fetal development has more than one start point, and thus we attempted to first resolve end points in the data (Figure S2A). The endpoints as defined here reflect cells that have reached a point in differentiation where they do not function as progenitors for other cells in the dataset as predicted by the manifold but might perhaps not yet constitute mature cell types at this PCW 8-10. To achieve this, we calculated the diffusion pseudotime for all clusters, and proceeded to walk along the K-nearest neighbour (KNN) graph between all clusters (see Methods). Using this approach, we could observe that some paths are rarely traversed, except in the case where the cluster is an end point. Thus, we designated clusters that were rarely traversed as end points and used the cell cycle score as obtained from Seurat (Stuart et al., 2019) to identify highly cycling populations, likely to be wrongly assigned end points which were subsequently removed. To obtain lineage information, we proceeded to walk back from all clusters across the manifold (see Methods). Lineage scores were then calculated, and lineage membership was determined by calculating the percent of lineage influence over all cells compared to all lineages (Figure S2B). We calculated the overall position of each cell in development as the sum of all the lineage scores and defined the differentiation score as the highest lineage membership value for each cell. We validated this approach by using a pancreas single cell RNA-seq dataset (Bergen et al., 2019; Lange et al., 2020) where trajectories have been previously defined (Figure S2C-F). Interestingly, the differentiation score recapitulated the endpoints although they are calculated using two different methods (graph walk and back diffusion), strengthening the confidence in end point prediction. Using the differentiation score, we defined undifferentiated cells as those obtaining a low score and differentiated cells as the cells that obtain a high score (Figure 2B, C). The obtained differentiation score for all cells anti-correlated with cycling and radial glial expressed genes such as *HMGN2*, *PCLAF*, *TYMS, H2AX, HMGN2, MAD2L1, ZFP36L1*, *GLI3*, and others, and positively correlated with *STMN2, MLLT11, TMSB10, TUBB2A*, *NEUROD6*, *DCX*, *MAPT*, *BCL11A*, *SYT1*, and others (Pearsons R, Fisher test, FDR 5%). To obtain more functional insights in the two genesets, we performed a pathway enrichment where we found the anticorrelating gene set to be enriched in pathways relating to cell-cycle checkpoints, DNA methylation, RNA polymerase I promoter opening, telomere maintenance and more (Figures S3B-G). Conversely the gene set positively correlating with differentiation was enriched in pathways relating to axon pathfinding by SLIT and ROBO, acetylcholine-, serotonin-, norepinephrine-, and glutamate neurotransmitter release, NMDA receptor activation and postsynaptic events, long-term potentiation, and others (Figures S3B-G). Thus, the differentiation score correlated with well-known pathways active in both stem-cell maintenance, and neural differentiation, suggesting the score captured a general progression from undifferentiated progenitors towards maturing clusters of cells in the developing forebrain.

We then proceeded to infer the putative OL lineage and calculated the lineage membership score. To approximate the accuracy of the OL lineage score, we compared it to the joint expression profile of the major known OL lineage markers *OLIG2, OLIG1, NKX2-1, NKX2-2, and SOX10* (Figure 2A). This showed moderate agreement as measured by Pearson R of 0.59 between the gene module total counts, and the calculated lineage membership (Figure 2A). We next focused on resolving transitions that occur within the putative OL lineage. We designated pseudotime as the sum of all the end point lineage scores over all cells. To select lineage associated genes, we ranked genes using KL divergence (see Methods), we selected the top 1000 genes scoring high for the lineage and modelled the expression using a generalized additive model (GAM) implemented by the mgcv R package (Wood et al., 2016) to smooth the expression along pseudotime. Significant genes were selected using Spearman correlation against the lineage membership score (Spearman Rho, Fisher test, FDR 1%) and the smoothed GAM predictions were min max normalized and ordered based on their maximum predicted expression point along the pseudotime (Figure 2D, E, Figure S3A).

We observed that the pseudotime determined early populations within the cells with estimated OL lineage potential showed high expression of patterning factors associated with the lateral and medial ganglionic eminence (LGE and MGE, respectively) such as *DLX1*, *DLX2*, *GSX1*, *GSX2*, and *NKX2-1* (Figure 2D), within the glioblasts clusters 7 and 15 (Figure 2G). Furthermore, our ranking and subsequent correlation test associated *NR2F1/2* (also known as *COUP TF 1/2*) with the OL lineage (Figure 2D). *NR2F1* and *NR2F2* are found to be necessary for both neural progenitor specification and glial commitment in CNS derived neural progenitor cells (NPCs) (Naka et al., 2008), and consistently expressed in the predicted lineage cells, possibly indicating that the radial glial populations are already glial potent at this early developmental stage in human. Interestingly, we also observed another radial glial sub-population within clusters 1 and 11 with estimated OL lineage potential, that did not express (MGE) patterning factors, but instead expressed *HOPX*, and to a lesser degree *LHX5*, *SP5* and *GATA3*, which we defined as outer radial glial cells (Figures 2D and 2F, Figure S3A).

The predicted OPC trajectory suggests that both MGE and outer radial glia progenitor groups progress through a stage of *ASCL1* upregulation combined with *EGFR*, and *HES6* expression after which the expression of key OL related genes occurs such as *NKX2-2*, *SOX10*, *PDGFRA*, *EDNRB*, *OMG*, *APOD*, and upregulation of existing expression of *OLIG2* among many other genes (Figure 2D,F-G, Figure S1D, S3A). Moreover, to visualize regulatory changes, we min/max normalized the smoothed expression of both the unspliced and spliced transcripts and visualized the difference between the two for each gene as the residual. Here we could observe different timepoints along the trajectory up- or downregulating genes such as early downregulation of *DLX5* and subsequent upregulation of high mobility group proteins *HMGN2* and *HMGB2*, known for involvement in transcription and chromatin remodeling(Ugrinova et al., 2009). We then observed high residuals for *ASCL1*, followed by *SLIT2*, *TFDP2*, *WLS*, *MEST*, *NR2F2*, *EGFR*, *SFRP2*, and *ZFP36L1* and others, indicating upregulation of developmental morphogens, and transcription factors along the trajectory (Figure 2E, Figure S1D). Thus, our predicted OL trajectory overlaps with parts of clusters 12, 6, 4, 23, 11, 1, 7, and 15, marking them as OPC progenitor populations and suggesting that the differentiation towards OPCs originates from the MGE and outer radial glial regions expressing regional patterning factors such *NKX2-1*, *GSX1/2*, and *HOPX* respectively, involves several transcriptional transitions, most notably *EGFR, ZFP36L1*, and *HES6* expression, and thus involves multiple progenitor populations and intermediate states at the PCW 8-10 timepoint

### Molecular definition of pre-OPC to OPC transition in the first trimester human forebrain

Our results indicate that the predicted oligodendrocyte lineage travels through the pre-OPC sub-clusters with a glioblast identity (cluster 15). Therefore, we proceeded to compare the predicted trajectories of both the glioblast population and the OPCs in more detail, to determine predicted overlap as determined by the lineage membership scores and which genes are correlated with either trajectory in a divergent manner. We attributed cells to each lineage by a low cut off value for the lineage membership score of 0.01 and performed Pearson correlation within these lineage cells, correlating for both the gliogenic membership score and the OPC membership score. We plotted the lineage scores of each end point to visualize the lineage structure. After an initial root structure, cells seemed to shift towards either fate indicating a branching structure (Figure S4A). We correlated expression with the lineage memberships of the OPC and glioblast lineages respectively (Spearman rho, cut off < 0.1, FDR 5%) (Figure S4B, C). Top positively correlating genes within the Glioblast and the OPC lineage biasing towards a OPC lineage fate included the patterning genes *DLX5*, *DLX1* and the proto-oncogene *MLLT11* known to act as a co-factor for *TCF4*. Other highly correlating genes included *ELAVL4*, a broadly acting RNA-binding protein involved in neural differentiation, *INSM1* a transcription factor necessary for the formation of the SVZ and delamination of radial glia (Tavano et al., 2018), also implicated in glioblastoma (Rosenbaum et al., 2015), possibly indicating that the glioblast lineage is not enriched for delaminated cells. Furthermore, *SOX4*, and *SOX11* were highly correlated with OPC lineage membership and were downregulated upon reaching the OPC state. (Figure S4B, C). *SOX4* and *SOX11* are known inhibitors of further OL differentiation, whereas *SOX10* is only expressed upon attainment of OPC fate (Claus Stolt et al., 2002; Potzner et al., 2007; Stolt and Wegner, 2010; Wittstatt et al., 2019). Additionally, we found expression of *DCX*, and *STMN2* (both microtubule associated proteins). Conversely, genes correlating with glioblast fate were broadly shared with OPC fate and captured more general gliogenic and radial glial factors such as *TTYH1*, *HES1*, *TSC22D4* and others, in accordance with a more undifferentiated state (Figure S4B, C).

### Regulatory networks underlying pre-OPC to OPC transition

To identify transcription factors regulating the transition of cells towards the OL lineage, we used SCENIC (Aibar et al., 2017), a framework to infer gene regulatory networks (GRNs) and transcription factor regulon activity using the GENIE3 algorithm (Huynh-Thu et al., 2010) (Figure 3A). We performed a Pearson correlation between the obtained regulon activity scores and the differentiation score over the entire dataset to obtain regulons that were significantly associated with differentiation scores (Pearson R, Fisher test, FDR 5%). Within the top significantly correlating regulons of the differentiation score were several known lineage regulators such as *KLF7*, a known regulator of differentiation in neuronal, cardiac and corneal epithelial differentiation, *SOX11*, *SOX4*, *TCF7L2*, *LHX9* amongst others (Figure S4D). Regulons correlating with undifferentiated states were mainly related to the cell cycle, such as *E2F7*, *E2F2*, and *TFDP2* which binds E2F family members, also highly correlating were nuclear receptor *NR2E1*, and *NFE2L2* a cytoprotective regulator (Figure S4E). Reactome pathway enrichment analysis of enriched regulons indicated enrichment of regulators of the cell cycle including *TP53*, regulators of senescence, regulators involved in SMAD signalling, and regulators of pre-NOTCH signalling, among other pathways (Figure S4F, G). Pathways enriched for the differentiation associated regulons included *TFAP2* regulatory pathway, regulators of beta-catenin, TCF complex, and HOX regulated hindbrain development, indicating possible contaminating or migrating populations in our sampled cells (Figure S4F, G). To infer regulons that play a role in OL differentiation we correlated the OL lineage scores with the regulon activity scores to assess regulons that were significantly associated with the lineage (Pearson R, Fisher test, FDR 5%, Figure 3B, C). The highest correlating motif activities were well known regulators of OL lineage specification and maintenance, such as *OLIG2*, *SOX10*, *SOX8*, *NKX2-2*, and *SOX6*, other highly correlating motif scores were patterning factors *GSX2*, *GSX1* and *DLX1*, and motifs for other factors such as *NKX2-3*, *E2F4*, *FOXB1*, *STAT3, SALL3*, and *SOX2* (Figure 3B, C). E2F1 has been implicated in cell-cycle control and E2F4 in repression of targets of E2F1, and has been found to be important in regulating cell cycle exit (Magri et al., 2014). Both E2F1 and E2F4 are expressed in the OPC lineage, indicating a possible role for cell cycle exit in the differentiation trajectory towards OPCs. Furthermore, FOXB1 has been revealed to be involved in inhibiting the cell cycle during oligodendrogenesis, but promoting differentiation in the developing forebrain (Zhang et al., 2017) (Figure 3 A, B). We calculated enrichment of regulon activity across the dataset, we observed transcription factor grouping broadly in accordance with the clustering, notably we observed distinct separation of cells based on patterning genes such as *DLX1/2/5/6* and overlapping *LHX6* and *SOX6*. Interestingly, although both *GSX1* and *GSX2* motif activity was predicted to be active throughout the OL lineage, GSX1 appeared to have a higher predicted activity in the glioblast cluster (cluster 15) compared to GSX2 (Figure 3A). Additionally, radial glial populations grouped together enriched for cell cycle regulatory factors such as *E2F2*, *E2F7*, *TGIF1*, *STAT1*.

**Figure 3.**
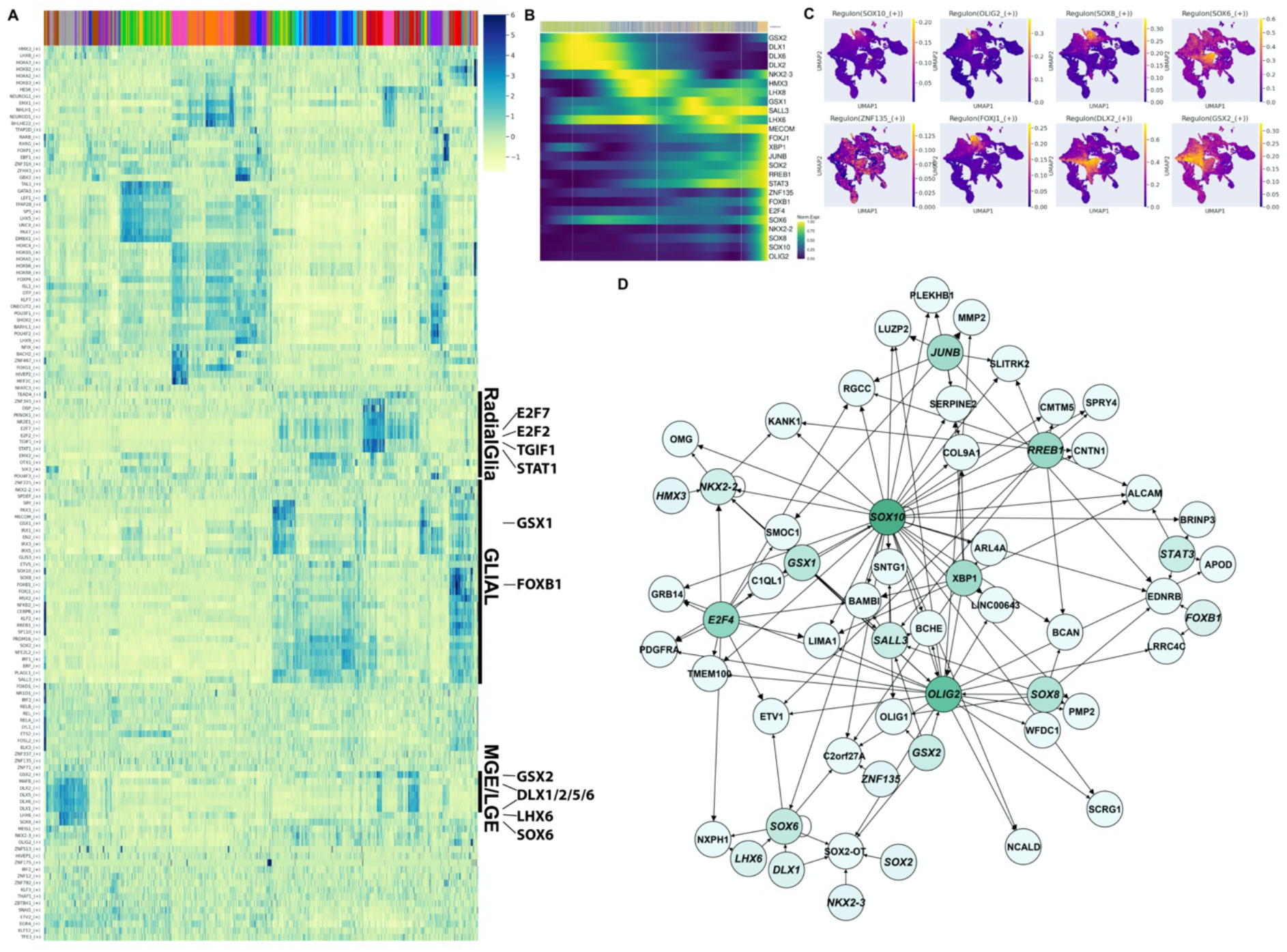
Transcriptional network connects early and late regulon activity along the OL lineage. A) Heatmap of enriched regulons for all cells in the dataset (Z-score of regulon activity). B) Top 25 regulons, estimated to be lineage drivers due to correlation to the OL lineage membership score. C) Selection of regulons activity profiles related to the OL lineage. D) Regulon network of top 20 regulons and top 100 genes associated to the lineage.

To assess the predicted regulatory interactions from the GRN generated using SCENIC, we selected the top 20 regulons and top 100 genes associated to the lineage (Pearson R, Fisher test, FDR 5%) and visualized the connected network after filtering for single edges, so that any target shares at least two transcription factors in the network. As expected, the regulon network indicated a strong positive feedback exists for *OLIG2*, *SOX6*, and *SOX10*, and strong interconnectedness for the major OL lineage transcription factors. Additionally, both *GSX1* and *GSX2* were predicted to regulate *OLIG2*, and *SALL3*, whereas *GSX2* also showed a direct activation of *OLIG1* (Figure 3D). This suggests that the transcription factors *GSX1/2* might play a vital role in early forebrain OPC development. Interestingly, *SALL3* was predicted to be mainly targeted by factors that were active in an early timepoint of the predicted trajectory (Figure 3B, D). Since *SALL3* targets the major OL lineage factors *OLIG1*, *OLIG2*, and *NKX2-2* (Figure 3D), it might play an important role for OL lineage initiation during human forebrain development.

### Regulons along the preOPC to OPC transition involve NOTCH signaling

We then turned to identifying genes that might be involved in early budding from the glioblast lineage towards OPC fate. To achieve this, we chose a more stringent cut off for glioblast lineage membership (cut off is 0.5, where more than half of the cells membership belongs to the glioblast lineage) and used Pearson correlation on the OPC lineage membership score within the glioblast lineage (> 0.1 Pearson’s R, 0.05 FDR). We found highly significant genes which we categorized into a OPC UP and DOWN module, with the score reflecting the mean expression of the genes in the module, showing a clear increase in expression when the OPC lineage score increased (Figure 4A). We found many genes that were previously predicted in the OPC lineage analysis. Highly correlating genes included *EGFR*, *DLL1/3*, *HES6*, *HNRNPA1*-(P48), *ASCL1*, *GLCCI1*, *OLIG1*, *OLIG2*, *CD24*, *GADD45G*, *TFDP2*, *ZEB1*, *SOX4*, *INSM1* and others (Figure 4B, Figure S4J). We observed genes expressed in the pre-OPC to OPC transition that seemed to be shared with other neuroblast populations. The shared genes included *ZEB1* which is known to be associated with the epithelial to mesenchymal transition in cancer, and known the be associated with *DLL1*, *DLL3*, *NOTCH*, *MAML1/2/3*, *JAG1*, *NOTCH1*, leading to attenuation of NOTCH signaling (Brabletz et al., 2011). The most strongly correlated genes with early OPC differentiation from the pre-OPC state involves both EGFR and NOTCH signaling, through the EGFR receptor, and NOTCH signaling ligands *DLL1, DLL3, MAML1/2/3,* and *NOTCH1* expressed in the OPC trajectory (Figure S4J). Furthermore, *MFNG* a modulator of *NOTCH1*, enhancing the interaction with *DLL1* is also upregulated in this transition, implicating NOTCH signaling as a major component of the pre-OPC to OPC transition (Brabletz et al., 2011).

**Figure 4.**
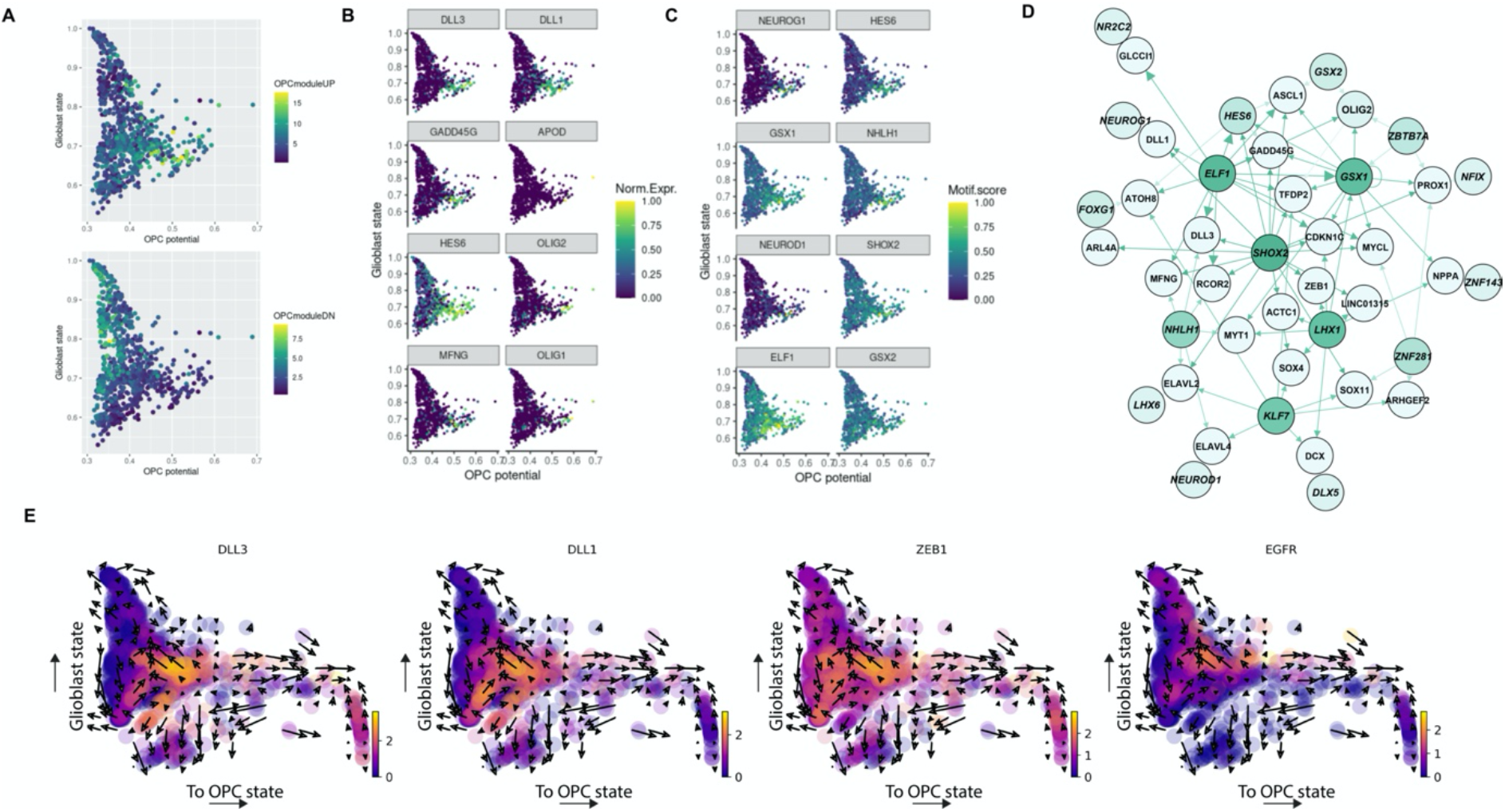
Regulons along the preOPC to OPC transition involves NOTCH signaling. . A) Module scores for the top genes correlating with OPC “budding” from the glioblast state. B, C) Top 8 correlating genes and regulons with OPC “budding” from the glioblast lineages. D) Regulon network of top 20 regulons and top 100 genes associated to the “budding” from the glioblast lineage. E) Velocity vector field showing lineage score towards glioblast state and OPC state respectively, illustrating the relatively large RNA velocity change accompanied with the initiation of the OPC state progression. Colored by normalized expression of each respective gene.

To infer the regulon activity profiles correlated with the switch to the OPC state, we generated a network with the top 20 regulons and top 100 target genes, we filtered targets that had only a single predicted transcription factor associated to them and visualized the network (Figure 4C, D, and Figure S4J). Highly correlating regulons with OPC lineage membership in the glioblast lineage included *NEUROG1*, *HES6*, *GSX1*, *NHLH1*, *NEUROD1*, *SHOX2*, *ELF1*, and *GSX2*. The constructed regulon network indicated that major connected predicted drivers of the network by out-degree were *ELF1*, *SHOX2*, *GSX1*, *LHX1*, *KLF7*, *NHLH1*. SHOX2 was the most central to the network and seemed to be expressed during the transition (Figure 4C, D and Figure S4H-J). However, SHOX2 has a role supporting OLIG2 in the pMN formation suppressing motor neuron fate (Hochstim, Christian John, 2009). The network did not share many regulons with the OL lineage network discussed previously, indicating that these processes might be driving different developmental processes. Additionally, many of the targets of the preOPC to OPC transition were expressed in other differentiation processes in the dataset indicating a more general network that might be adapted by patterning factors such as OLIG2 and GSX1 in this network.

We next investigated the predicted RNA velocity vector field associated with the OL and glioblast lineage (lineage membership cutoff> 0.1) and observed that most genes were expressed right before or during major transcriptional shifts occurred towards the OPC state. We observed *EGFR*, *DLL1/3*, and *ZEB1* upregulate expression right before the shift to the OPC state occurs (Figure 4E). Interestingly, Jagged1 and Contactin1 have been found to have inhibitory and promoting functions in OPC differentiation (Hu et al., 2003; Wang et al., 1998), which we could observe in our data, where we see JAG1 downregulated upon OPC differentiation, witch a marked upregulation of CNTN1 whilst we observe maintained expression of the NOTCH1 receptor during differentiation (Figure S4J). Furthermore, we could observe upregulation of OPC marker genes such as PDGFRA, and APOD, but also, interestingly expression of DCX, previously found in OPCs (Boulanger and Messier, 2017) (Figure S4J). EGFR^+^ DLL1^+^ transit amplifying cells have been shown to asymmetrically divide leading to differential inheritance of receptors during VZ/SVZ division. Additionally, it has been shown that the EGFR^+^DLL1^+^ transit amplifying cells require DLL1 for maintaining and promoting quiescence, thereby promoting their own quiescence (Kawaguchi et al., 2013). A recent study on later stage human development also identified asymmetric division from an EGFR+ progenitor population giving rise to OPCs (Huang et al., 2020). Collectively, this would suggest a role for EGFR and DLL1 regulation as well as NOTCH signaling in the asymmetric division upon OPC fate acquisition and suggests that the cell cycle arrest genes, *GADD45G*, and *TP53* play a role in obtaining a quiescent pool of oligodendrocyte progenitors in the SVZ.

### Chromatin accessibility compatible with oligodendrogenesis in the first trimester human forebrain

Our single-cell transcriptomics analysis indicates that oligodendrogenesis already occurs between PCW 8-10. For such a OL transcriptional programme to be executed, chromatin at regulatory regions of the genes involved need to be in open, accessible states. To investigate whether this was indeed the case, we performed single-cell assay for transposase accessible chromatin followed by sequencing (scATAC-seq) (10x Genomics) on human forebrain in the first trimester, at PCW 8.5, 9.5 and 11 (Figure 5A and 5B). Using the scRNA-seq annotation (Figure 1B) and label transfer, we first mapped scATAC activity scores to gene expression. Seurat finds gene anchors between both modalities in low dimensional space and builds a cell type classifier based on the reference dataset. For every cell, we obtain a prediction score for each cell type, which can be interpreted as a similarity score (Stuart et al., 2019). This approach allowed the identification of 18 out of 32 clusters found in the scRNA-seq, with prediction scores higher than 0.7 (Figure 5A and Figure S5B). Most cells have a high prediction score for a specific cell type and a low for all others (Fig. S5B). Among those, we found 51 cells. similar to OPCs and 562 similar to pre-OPCs. While OPCs clustered close to pre-OPCs, we obtained a prediction score higher than 0.6 for OPCs and lower than 0.3 for pre-OPCs, suggesting a distinct chromatin accessibility profile (Figure S5B). We identified 207 genes with increased activity (padj < 0.01) between OPCs and pre-OPCs (Supplementary Table 2). In concordance with the scRNA-seq, genes such as *SOX10, BRINP1* and *CMTM5* have highly increased activity scores in OPCs (Figure 6A). Reversibly, pre-OPCs presented higher chromatin accessibility in *CRB2, SOX1-OT* and *A2ML1* (Figure 6A). We also found that the chromatin at regulatory regions controlling some genes involved in the OPC specification, such as *LUZP2, PDGFRA* and *ETV1* were already open in pre-OPCs (Figure 6A), despite their reduced expression (Figure 6C), suggesting chromatin priming prior to OPC specification. In addition, we found enriched motifs of transcription factors such as OLIG1, OLIG2 and EPAS1 in OPCs when compared to pre-OPCs (Figure 6B). In contrast, we found that JUNB, KLF4 and SOX10 have enriched motifs only in a portion of the OPCs (Figure 6B). Thus, our results indicate that the human forebrain contains at the first trimester subsets of cells with a chromatin state compatible with oligodendrogenesis.

**Figure 5.**
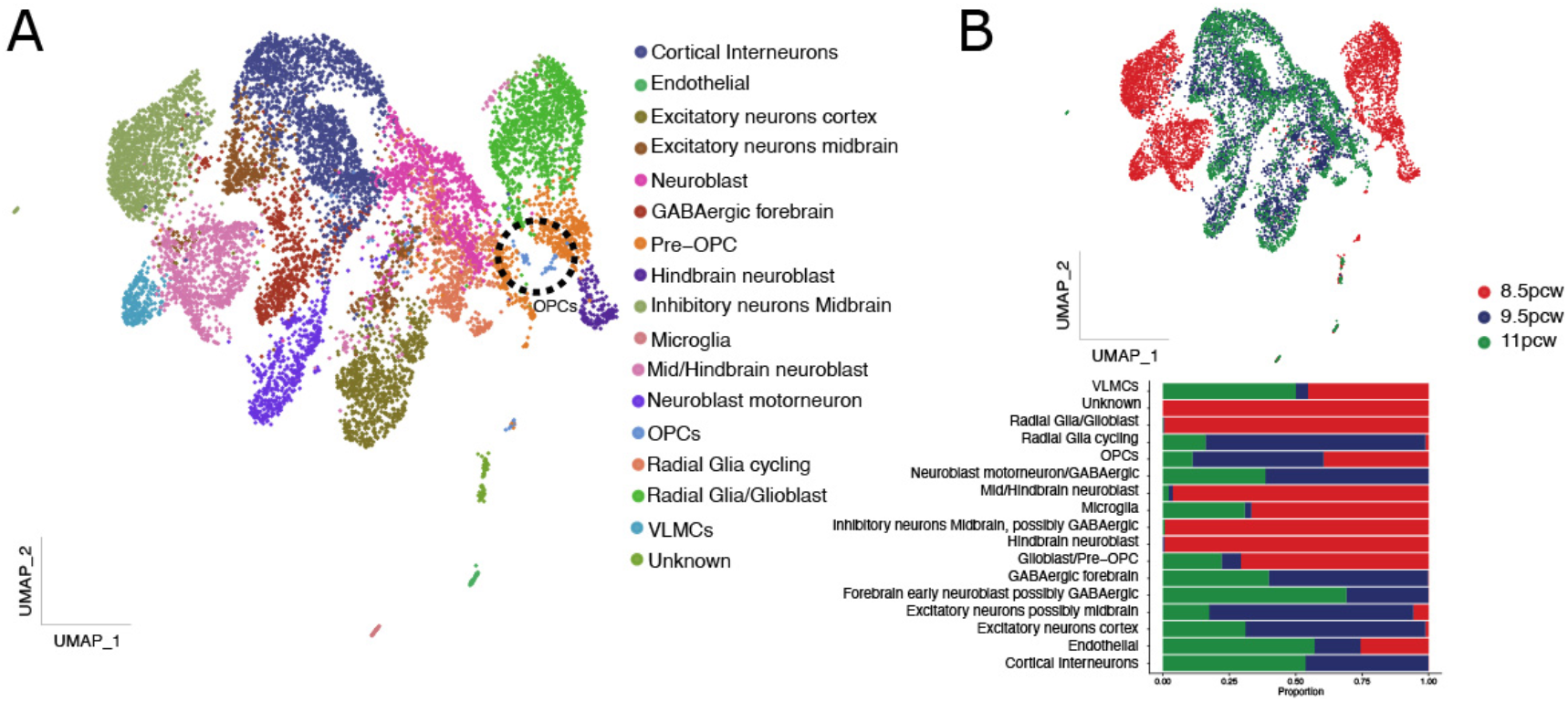
Chromatin accessibility on the OPC lineage in first trimester human forebrain. A) UMAP of 10487 cells distributed in 17 clusters annotated via label transfer with the scRNA-seq as reference. B) Cell distribution according to post-conceptional week age in UMAP representation (upper) and its proportion in each cluster.

**Figure 6.**
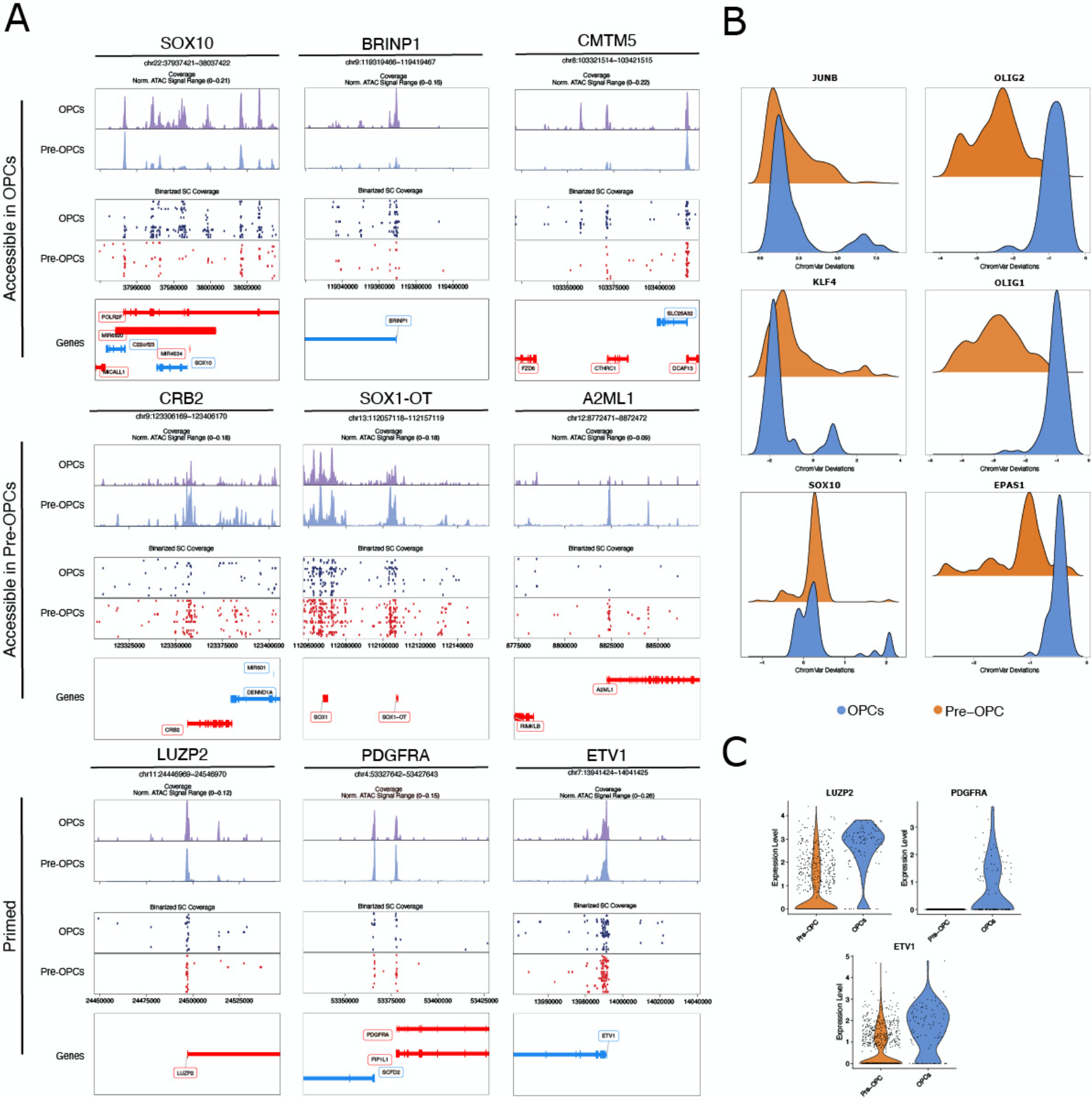
Accessible genes with chromatin acessibility in OPCs and preOPCs. A) Track plots of genes differentially accessible in OPCs (upper), pre-OPCs (middle) or primed in pre-OPCS (lower). The first row in each plot represents normalized pseudo-bulk coverage. Second row represents a sample of 30 cells from each cluster and its binary signal. Last row depicts the locus. B) Density plot of ChromVar motif deviation score in OPCS and pre-OPCS. C) Expression of genes with chromatin accessibility in both OPCs and Pre-OPCs.

### Spatial ISS analysis defines the ventral forebrain as a site for early human oligodendrogenesis

Our single cell omics analysis between PCW 8-11 indicates that oligodendrogenesis occurs in the human forebrain, with two likely origins, outer radial glia and MGE radial glia. To determine the exact spatial location of the first waves of OL lineage cells, we performed In Situ Sequencing (Gyllborg et al., 2020; Ke et al., 2013) on an entire PCW 8 forebrain, with 10μm thick coronal sections covering both ventral and dorsal regions and spanning three regions in the anterior-posterior axis (Figure 7A-E, S6 and S7). In particular, we applied hybridization-based In Situ Sequencing (HybISS) (Gyllborg et al., 2020) that has an increased signal-to-noise ratio and performs well for in situ mapping of human cell types (Langseth et al., 2021). We targeted 50 genes which characterize subsets of the populations identified_by scRNA-seq and scATAC-Seq (Figure 7D, E and S7A-D) and used pciSeq (Qian et al., 2020) for cellular assignment. While HybISS has single-cell resolution, we noted that the human forebrain at this stage has a high cellular density (Figure S7D)(Nowakowski et al., 2017; Pollen et al., 2015), and thus applied a grid-based approach for cell assignments. As expected, we observed radial glia (cluster 11) lining the entire breadth of ventricles, neural progenitors (cluster 14) in the sub-ventricular zone, while early inhibitory (cluster 28) and excitatory neurons (clusters 9, 16, 28) at the parenchyma, with the latter clusters positioned in the periphery of the MGE (Figure 7A, B and S6). Radial glia clusters 4 and 18, and partial 11 were enriched in the ventral forebrain, but interestingly in the outer subventricular zone (Figure 7B) (Nowakowski et al., 2017; Pollen et al., 2015). Notably, *HOPX,* a marker of outer radial glia at the second trimester (Nowakowski et al., 2017; Pollen et al., 2015), was not expressed as much in these outer layers but rather at the ventricles (Figure 7E and Figure S7), suggesting that the outer radial glia at the first trimester might have different properties. We also identified neural progenitor populations (clusters 6, 17) and radial glia (cluster 21) present in the parenchyma, in areas enriched with differentiated excitatory neurons (Figure 7B and Figure S6). Glioblasts (cluster 15), which include pre-OPCs, lined the ventricles in a layer adjacent to radial glial cells (cluster 11). They were also assigned to the meninges and choroid plexus (Figure 7A, B and S6), which is most likely due to co-expression of some of the genes characterizing glioblasts by populations in these tissues. OPCs (cluster 26) were observed in ventral regions, namely in the medial ganglionic eminence (Figure 7B and S6). Interestingly, OPCs appeared to have already migrated from their sites of origin (outer subventricular zone and ventricular zone) and were observed in areas where excitatory neurons reside (Figure 7B and S6).

**Figure 7.**
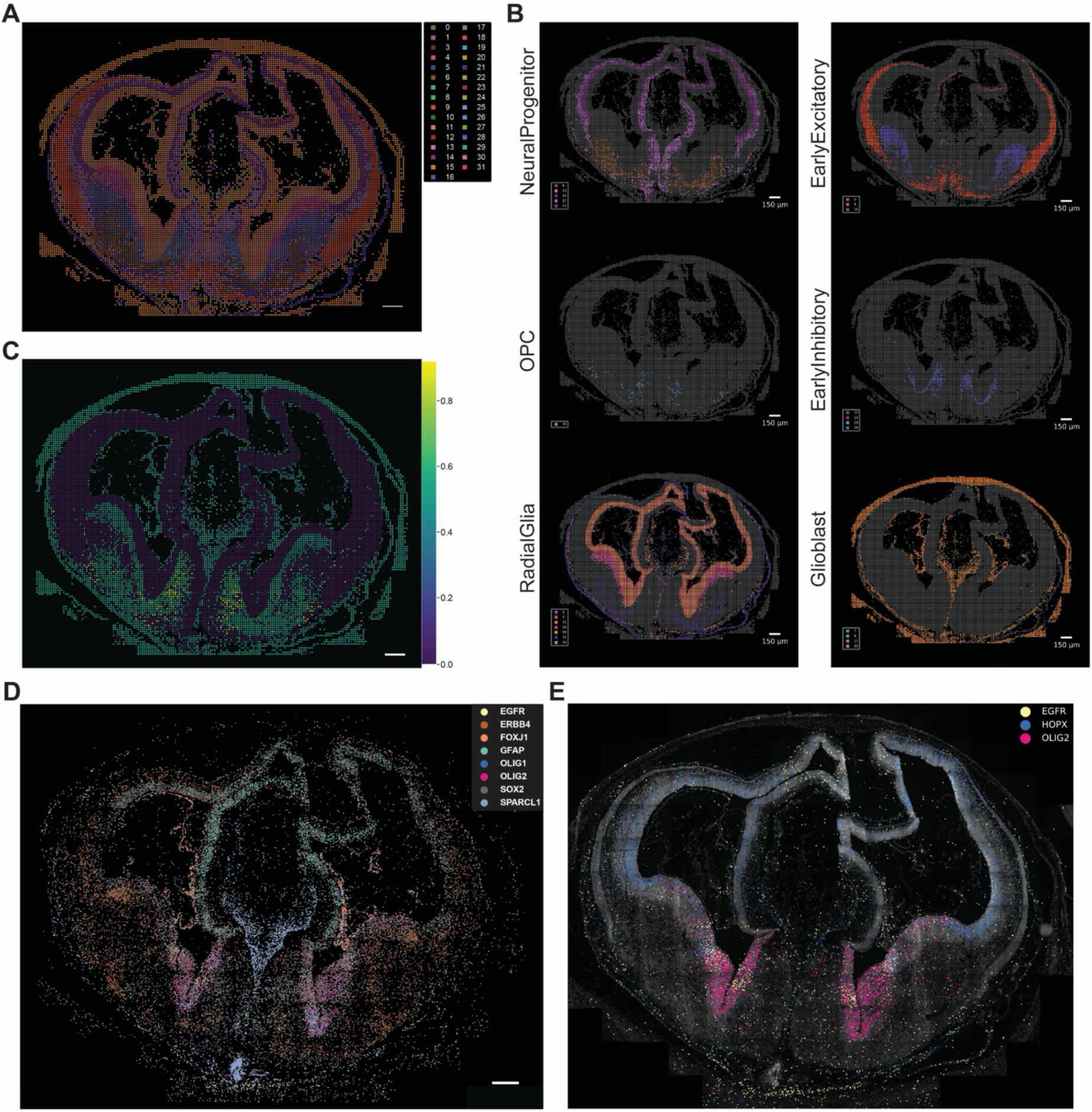
Spatial transcriptomics of PCW8 brain highlights the ventral forebrain as a region of early oligodendrogenesis. A) PciSeq HyISS showing all clusters from Figure 1B mapped to spatial coordinated using probabilistic mapping. B) Individual clusters grouped as shown in Figure 1C. C) The average OPC potential by cluster translated to spatial coordinates using the cluster assignment as in A (softscaled potential^0.1^ for visualization). D, E) Spot recordings of probes referring to a selection of genes showing DAPI staining in white.

We then investigated the locations where cells with higher estimated OL lineage potential (Figure 2A) were present by averaging the OL lineage potential for each cluster and visualizing the average cluster potential onto the spatial mapping (softscaled (potential0.1) for visualization purposes). As predicted, cells representing clusters with highest average potential were absent in dorsal regions, and were instead restricted to ventral domains, away from ventricles, suggesting migration. We noticed an increase in OPC potential near sites of EGFR expression (Figure 7C-E). In addition, only one site of OPC potential increase was matched with HOPX expression (Figure 7C, E), suggesting that at the PCW 8 timepoint not all OPCs are generated from HOPX+ regions. Other cells with much lower potential were present in the first trimester outer radial glia region, and also at the ventricles, where glioblasts are located, reflecting the partial overlap with the OL lineage resulting in much lower average potential values (Figure 7C). Future studies, perhaps with finer clustering and a bigger probe set could give a finer grained picture. Regardless, our spatial HybISS analysis confirms oligodendrogenesis in the human first trimester, from radial glial cells in the outer subventricular zone and glioblasts from the MGE.

## Discussion

In this study, we performed an extensive analysis of the identity of neural progenitors at the human forebrain at the first trimester, at PCW 8-11. Our results indicate that the oligodendrocyte lineage is specified already at this early stage of human gestation, several weeks from what was previously thought. Interestingly, oligodendrogenesis is also observed in the human first trimester spinal cord (Erik Sundström, personal communication, (Marklund et al., 2014; Rayon et al., 2021)). These findings suggest that human oligodendrogenesis occurs in several developmental waves, as previously described in mouse forebrain (Kessaris et al., 2006) and spinal cord (Fogarty et al., 2005). Interestingly, these waves in mouse undergo a process of transcriptional convergence, giving rise to OPCs with equivalent properties at post-natal stages (Marques et al., 2018; Zeisel et al., 2018), with similar potential regarding the capacity to differentiate into distinct mature OLs (Floriddia et al., 2020). This could also apply for the human OPC developmental waves, although further investigation will be required to determine if that is indeed the case, and what would be the biological functional of such waves.

Velocity back-tracing revealed several states along a possible trajectory of OPCs, which seem to be mainly NKX2-1, GSX1, and GSX2 restricted. Expression withinthese clusters indicates a departure from the VZ into the SVZ or beyond, perhaps becoming intermediate progenitors or transit amplifying cells. Our differential expression analysis between preOPC/glioblast population (cluster 15) and OPCs show great differences indicating a considerable transcriptional overhaul in a seemingly short developmental window. Our data also suggests two distinct EGFR+ populations as sources of OPC. As Kriegstein and colleagues have previously shown (Nowakowski et al., 2017; Pollen et al., 2015) that outer radial glia at the second trimester also has this capacity. Our results indicate that at the first trimester, both radial glia and a HOPX-negative outer radial glia population also have the potential to become OPCs. Thus, in this aspect human oligodendrogenesis differs from mouse, where the first wave is restricted to the MGE (Kessaris et al., 2006), while in human at least two distinct domains of origin occur in the first trimester.

Further analysis of regulatory network controlling the transition between pre-OPCs to OPCs highlights canonical OL transcription factors such as SOX10, OLIG2, SOX8, NKX2.1, among others but also others that have not been previously involved in this process, such as SALL3, LUZP2, NCALD, ETV1, MITF, TRAF4 and DLX genes. The expression of DLX genes in the populations of origin seem to contradict conventional transcriptional programs where oligodendrocyte fate is repressed by DLX genes. Although it has been reported that DLX positive progenitor populations can generate OPCs (He et al., 2001; Kriegstein and Alvarez-Buylla, 2009), it is not clear if that is the case in our captured populations, and additionally, we observe downregulation of DLX genes before upregulation of *OLIG2* occurs. However, DLX5 expression is still observed in some OPCs indicating that DLX expression might repress OL fate, but not completely.

Our results also place NOTCH signaling central to OPC specification. NOTCH signaling had previously been shown to either prevent (Genoud et al., 2002; Hammond et al., 2014; Wang et al., 2017, 1998) or promote (Hu et al., 2003) OPC differentiation in mouse. Mouse and zebrafish OPC specification is driven by Notch signaling (Grandbarbe et al., 2003; Kim et al., 2008; Park, 2005) and our data indicates that this most likely is also the case in human. Future investigations will determine if the newly identified transcription factors and NOTCH signaling indeed are essential for these early transitions during oligodendrogenesis, and if they can be used to improve the reprogramming of pluripotent cells or differentiated cells into OPCs.

## Acknowledgements

We would like to thank Tony Jimenez-Beristain for assistance and Matthew Speir for setting up the UCSC Cell Browsers for the datasets. We acknowledge support from the Eukaryotic Single Cell Genomics (ESCG) facility in Stockholm funded by Science for Life Laboratory, KI Core and StratRegen, the National Genomics Infrastructure in Stockholm funded by Science for Life Laboratory, the Knut and Alice Wallenberg Foundation and the Swedish Research Council, and Swedish National Infrastructure for Computing /Uppsala Multidisciplinary Center for Advanced Computational Science for assistance with massively parallel sequencing and access to the UPPMAX computational infrastructure. Work in MN’s research group was supported by Chan Zuckerberg Initiative, an advised fund of Silicon Valley Community Foundation; Erling-Persson Family Foundation (A human developmental cell atlas); Knut and Alice Wallenberg Foundation (KAW 2018.0172); Swedish Research Council [2019-01238]: This project has been made possible in part by a grant from the Chan Zuckerberg Foundation (Seed Networks “Oligodendroglia heterogeneity in the human brain”, G.C.-B as grant coordinator). MN and GCB were co-funded by a grant from the Strategic Research Programme in Neuroscience (StratNeuro). Work in G.C.-B.’s research group was supported by the Swedish Research Council (grant 2015-03558 and 2019-01360), the European Union (FP7/Marie Curie Integration Grant EPIOPC, Horizon 2020 Research and Innovation Programme/European Research Council Consolidator Grant EPIScOPE, grant agreement number 681893), the Swedish Brain Foundation (FO2017-0075 and FO2018-0162), the Swedish Cancer Society (Cancerfonden; 190394 Pj), Knut and Alice Wallenberg Foundation (grants 2019-0107 and 2019-0089), The Swedish Society for Medical Research (SSMF, grant JUB2019), Olav Thon Foundation, Ming Wai Lau Centre for Reparative Medicine and Karolinska Institutet.

## Code and Data availability

The raw data is in the process of being deposited in European Genome-phenome Archive (EGA) (https://ega-archive.org). A link will be made available when ready. Code and R-objects of the data for the scRNA-Seq and snATAC-Seq are deposited at https://github.com/Castelo-Branco-lab/humandevOLG. For the HybISS, code can be found at https://github.com/Moldia/iss_starfish/ and https://github.com/acycliq/pciSeq. UCSC Cell Browsers (Speir et al., 2021) for visualization of the scRNA-seq and snATAC-Seq are available at https://human-forebraindev.cells.ucsc.edu.

## Author Contributions

DvB and GCB conceived and designed the study, DvB performed scRNA-seq and data analysis, FP performed snATAC-seq (with MK and MM) and data analysis, PK performed sample preparation and imaging for HybISS and IHC (not shown). HL performed sample preparation, library preparation, probe hybridizations and imaging for HybISS. CML performed image analysis, spot detection, gene calling and cell mapping analysis of HybISS data. MMH performed cell mapping analysis and supervised the HybISS experiments. DvB wrote the manuscript together with FP and GCB, with input from all authors.

## Methods

### Human Tissue

Human first trimester forebrain tissue was obtained from elective abortions (8-11 weeks post-conception) with written informed consent of the pregnant woman and in accordance with the ethical permit given by the Swedish Ethical Review Authority (Stockholm, Sweden, reference no. 2007/1477-31/3 with amendments 2011-1101-32, 2013-564-32, 2016-589-31, 2018_2497_32), and the National Board of Health and Welfare. Human fetal forebrain tissue was collected and stored in hibernation media with the addition of GlutaMAX and B-27 supplements according to the manufacturer’s instructions (overnight, 4 °C, Hibernate-A, Thermo-Fisher). Tissue was then cut into small cubic pieces of approximately 1–2 mm length. Tissue was dissociated using a dissociation kit (Miltenyi, Neural Tissue Dissociation Kit (P)) according to the manufacturer’s instructions. In brief, tissue was prepared in the kit buffer containing 0.067 mM β-mercaptoethanol. After addition of enzyme mix 1 and 2, the tissue was mechanically dissociated using three increasingly smaller gauges of fire-polished Pasteur pipettes, pipetted 20, 15 and 10 times up and down, respectively. Finally, collected cells were stored on ice in PBS containing 1% BSA and immediately prepared for single-cell library preparation. Single-cell RNA sequencing was performed using the 10x Genomics Chromium V2 and v3 kit, following the manufacturer’s protocol, and sequenced on an Illumina Hiseq 2500.

For scATAC-Seq, the nuclei Isolation for 10x Genomics Single Cell ATAC Sequencing demonstrated protocol was used (CG000169, 10x Genomics, 2019), and sequenced on an Illumina Hiseq 2500.

### Data analysis, clustering and integration with Harmony

FAST-q data obtained from single-cell RNA-seq was processed using Kalisto (Bray et al., 2016). The reads were separated into unspliced and spliced counts. Quality control was performed on the spliced matrix and only the cells that passed the quality control were analyzed further. After quality control we obtained 25161 cells which were subjected to downstream analysis. Cells were normalized so that the total counts in the cells summed up to 1 and then multiplied by a factor of 10000 to avoid any numerical problems.

We then calculated features using a common support vector regression model on the coefficient of variation versus the magnitude of expression for each gene, and performed dimensional reduction on the obtained expression matrix, after which we used Harmony (Korsunsky et al., 2019) to integrate technical batch effects due to V2 and V3 kits of 10X, on 50 components.

### Reintegration of V2 and V3 datasets

We projected the V2 and V3 data into each PCA reduced space, and generated a JSD matrix for both PCA spaces, which were then summed to create a distance matrix in which we determined nearest neighbours. We then proceeded to integrate the 10 nearest neighbours of each V3 cell in the V2 data and calculated the correction matrix as performed in Seurat and MNN (Haghverdi et al., 2018; Stuart et al., 2019), integrating V3 with V2 data.

### Generating count matrix and RNA velocity diffusion maps

To generate a manifold respecting both the RNA velocity and transcriptional similarity-based distances between cells, we used the corrected PCA space from the integration between the V2 and V3 data as input for scVelo (Bergen et al., 2019) to obtain a transition matrix of cell velocities. We generated a diffusion map using the destiny/DPT R package (Haghverdi et al., 2016) on the integrated PCA space to obtain a transition matrix of transcriptomic similarities. Next, we performed canonical correlation analysis (CCA) on both these transition probability matrices (velocity and integrated PCA space) and used these components (40 components) as input to the final diffusion map to generate a manifold and transition matrix congruent with transcriptional and velocity distances that can capture non-linear relationships. However, the directionality of the RNA velocity transition matrix is lost.

### Path tracing and end point determination

Using the generated manifold diffusionmap, we then traced paths from all clusters to all clusters by using the calculated diffusion pseudotimes. To trace from all populations to all others, we iterated the following sequence, we started from the one population and took small steps (10 nearest neighbours) in the direction of the desired fate (the source of the cluster to target). By only selecting the cells in that step on the network that actually were closer to the desired fate (as measured by diffusion pseudotime), and continuing with the new (now closer to the target population) position, we continued stepping until we reached the destination point (diffusion pseudotime = 0). After completing a walk from cluster A to all other clusters, we mapped the number of times a path was traced over each cell. We did this for all clusters. We then calculated a z-score for each cluster set and set the arbitrary cutoff of >0.5 to belong to a pathtrace. We then binarized the path to 0 or 1 and smoothed the path values over the manifold by taking nearby neighbours and using a weighted mean to smooth over the 100 nearest neighbours we iterated the smoothening for a total of 5 times, using the weights as defined by the transition matrix of the diffusionmap.

We then calculated end points as normalizing the pathvalues for each cluster and calculating the proportion of contribution of each path to each cell. We then set clusters that obtained a contribution of 0.8 or higher as end **Figure 5**. points.

### KL divergence expression transformation to rank genes

We transformed the expression matrix by calculating the KL divergence for each cell to every other cell. By first converting the expression of each cell to a probability distribution by normalizing over all counts of the cell, forcing each cells expression to sum to 1. For each cell compared to every other cell, we calculated the KL vector as the following.

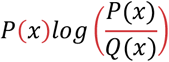

We then calculated the KL divergence as follows, again for each cell compared to every other cell.

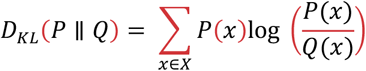

We then performed a matrix multiplication with the KL vectors of each cell multiplied by the KL divergence. After repeating the procedure for each cell, we obtain a matrix where each cells genes are ranked according to the relative distances between cells, we then normalized over the cells to have all gene contributions sum up to 1 for each cell. Over which we subsequently performed a comparison between cells to obtain a difference value to rank cells as follows. Where ClusterA and ClusterB are a vector for all the gene values belonging to each respective cluster. Where in this case ClusterB would be background (i.e. all cells not belonging to ClusterA)

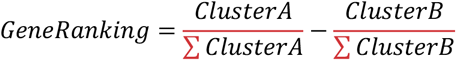

### Lineage backdiffusion and lineage membership assignment

To obtain lineage information, we proceeded to walk back from all clusters across the manifold. Here we used a similar approach as earlier used in CellRank (Lange et al., 2020), where we joined the Velocity transition matrix with the distance matrix based on counts. We combined both matrices in the following ratio 0.8 * backwards velocity ransition matrix + 0.2* transitionmatrix of diffusionmap based on counts. We then proceeded to walk back using weighted mean stats R package (R Development Core Team, 2010), by giving every clusters 50 most distant cells (as measured by the diffusion pseudotime from the path tracing steps) a value of 1, we then proceeded to diffuse back using weighted means with 1000 nearest neighbors, however as our kernel does not have values for all cells (K-NN cutoff in diffusion map) stetting a high threshold like 1000 neighbours would then automatically use the maximum cutoff in neighbors from both the transition matrix and the diffusionmap. We iterated this for 15 steps, and at the start of every diffusion step we added back the initial start values on top of the diffused values. Effectively feeding the diffusion to produce a gradient from each cluster diffusion back to startcells as measured using velocity. Lineage scores were then calculated, where we calculated lineage score as the KL divergence between lineages in the same manner as we did for cells in the ranking step (see KL divergence expression transformation to rank genes), but now for lineage information, so that lineages are now ranked according to the relative distances between cells, as dictated by the lineage scores. Lineage membership was determined by normalizing over the rows (lineages) and then normalizing over the columns (cells).

We calculated the overall position of each cell in development as the sum of all the lineage scores, and defined the differentiation score as the highest lineage membership value for each cell.

### Regulatory network

To calculate regulons we implemented the SCENIC pipeline (pySCENIC) (Aibar et al., 2017; Van de Sande et al., 2020) according to default settings.

### scATAC-seq analysis

Raw data was processed using CellRanger ATAC v1.2 platform. Reads were mapped to human reference genome GRCh38 provided by 10x Genomics. Most of the analysis was performed using ArchR v1.01 framework (https://doi.org/10.1038/s41588-021-00790-6) on R v4.1.0. In the quality control, cells were filtered based on the transcription start site (TSS) enrichment (>10) and the total number of fragments (>1000). After QC, ArchR used a 2kb genome widow to create a cell-by-bin matrix. At the same time, it also generates a gene activity matrix, calculating the number of reads on the gene body and distal regions that do not overlap with other gene bodies and normalizes by the distance from TSS. Dimensionality reduction was performed on the genomic bin space using an iterative (n=3) LSI (TF-IDF normalization followed by SVD). Clusters were obtained by Louvain clustering (resolution = 1). Following clustering, cells were annotated by label transfer with scRNA-seq using Seurat v4 (https://doi.org/10.1016/j.cell.2021.04.048). Clusters were annotated based on the proportion of each predicted cell type. Peak calling was performed by cluster with MACS2. Differential analysis (Wilcoxon) was performed on two levels: Gene activity and peaks. Transcription factors motif enrichment was performed using ChromVAR (doi: 10.1038/nmeth.4401.)

### Hybridization-based In Situ Sequencing (HybISS)

Human tissue at week 8 post conception was embedded in Tissue-Tek® O.C.T. (Sakura) and flash frozen in dry ice/ethanol bath, sectioned in 10 μm-thick cryosections and stored at −80°C until HybISS was applied. HybISS was performed as described by Gyllborg et al. (Gyllborg et al., 2020). Briefly, after fixation, sections were permeabilized with 0.1 M HCl and washed with PBS. A hydrophobic barrier was drawn along the tissue sections using an ImmEdge Hydrophobic Barrier PAP Pen (Vector Laboratories) to keep reagents localized over the tissue sections. cDNA synthesis was performed by reverse transcription overnight with reverse transcriptase (BLIRT), RNase inhibitor, and priming with random decamers. The next day, sections were post-fixed before padlock probe (PLP) hybridization and ligation at a final concentration of 10 nM/PLP, with Tth Ligase and RNaseH (BLIRT). This was performed at 37°C for 30 min and then moved to 45°C for 1.5 h. Sections were washed with PBS and RCA was performed with phi29 polymerase (Monserate) and Exonuclease I (Thermo Scientific) overnight at 30°C. Bridge-probes (10 nM) were hybridized at room temperature (RT) for 1 h in hybridization buffer (2XSSC, 20% formamide). This was followed by hybridization of readout detection probes (100 nM) and DAPI (Biotium) in hybridization buffer for 1h at RT. Sections were washed with PBS and mounted with SlowFade Gold Antifade Mountant (Thermo Fisher Scientific). After each imaging round, coverslips were removed and sections washed 5 times with 2XSSC and then bridge-probe/detection oligos were stripped with 65% formamide and 2XSSC for 30 min at 30°C. This was followed by 5 washes with 2XSSC. Now the next cycle of bridge-probes could be hybridized as above.

Imaging was performed with a Leica DMi8 epifluorescence microscope equipped with LED light source (Lumencor® SPECTRA X), sCMOS camera (Leica DFC9000GTC), and 20× objective (HC PL APO, 0.80). Each field-of-view (FOV) was imaged with 21 z-stack planes with 0.5 μm spacing and 10% overlap between FOVs.

### Image processing and decoding of HybISS data

After imaging, each FOV was maximum intensity projected to obtain a flattened two-dimensional image. Imaging data was then analyzed with in-house custom software that handles image processing and gene calling based on the python package Starfish. Each two-dimensional FOV was exported, and preprocessed including alignment between cycles and stitched together using the MIST algorithm. Stitching was followed by retiling to create smaller non-overlapping 6000×6000 pixel images that were then used for decoding. The decoding pipeline can be found on the Moldia GitHub page (**https://github.com/Moldia/iss_starfish/**). In short, the images were initially filtered (using the Filter module from Starfish) applying a white top hat filter with a masking radius of 15. The filtered images were subsequently normalized (using the Filter module from Starfish). Following the normalization, spots were detected using the FindSpots module from Starfish and decoded using PerRoundMaxCannel decoding.

### Cell type mapping

Probabilistic cell maps were created using probabilistic cell typing by in situ sequencing (pciSeq; **https://github.com/acycliq/pciSeq**, REF Qian et al., 2020). pciSeq assigns genes to cells and cells to cell types. The assignment is done using a probabilistic framework based on a single-cell RNA sequencing data. Due to the density of nuclei in the tissue, a compartment-based approach was employed in which each compartment was defined as 50×50 pixel grid (~16×16 μm).

## Legends for Supplementary Figures

**Figure S1.** A) Violin plot showing the distribution of the number of counts and number of features over all samples. B) Violin plot showing the distribution of the number of counts and number of features over all clusters. C) Barplot showing the distribution of age over the clusters. D) UMAP plots of selected genes. E) Volcano-plot showing differential expression results for preOPC compared to OPC. F) Volcano-plot showing differential expression results for preOPC compared to OPC but only annotated transcription factors and co-factors.

**Figure S2.** A) Left, UMAP showing the clustering performed for the lineage analysis after integration. Right, Endpoint assignment results of the lineage analysis. B) Lineage membership scores calculated for each cluster (softscaled (exp)0.1). C) UMAP of the benchmark test set as from the pancreatic dataset. D) Endpoint assignment results of the lineage analysis on the pancreatic dataset. E) Lineage membership scores for all clusters of the pancreatic dataset. F) Gene ordering and cell ordering towards the Alpha lineage in the pancreatic dataset.

**Figure S3.** A) Gene ordering and cell ordering towards the Oligodendrocyte lineage. B) Top 16 genes positively correlating with the differentiation score over the whole dataset (Pearson R, Fishertest, FDR 5%). C) Top 16 genes negatively correlating with the differentiation score over the whole dataset (Pearson R, Fishertest, FDR 5%). D) Reactome pathway analysis over the significantly positively correlating genes over the whole dataset (Pearson R, Fishertest, FDR 5%). E) Network view of results in D. F) Reactome pathway analysis over the significantly negatively correlating genes over the whole dataset (Pearson R, Fishertest, FDR 5%). G) Network view of results in F.

**Figure S4.** A) Custom layout of OPC lineage score on the x-axis and glioblast lineage score on the y-axis, showing shared cells between the lineages and a branching point towards the OPC state and glioblast state (arrows). B,C) Top 16 correlating genes for OPC (B) and glioblast (C) lineages (Pearson R, Fishertest, FDR 5%), D) Top 16 regulon scores positively correlating with the differentiation score over the whole dataset (Pearson R, Fishertest, FDR 5%). E) Top 16 regulon scores negatively correlating with the differentiation score over the whole dataset (Pearson R, Fishertest, FDR 5%). F) Reactome pathway analysis over the significantly correlating regulons over the whole dataset (Pearson R, Fishertest, FDR 5%), for both positive and negative correlation respectively. G) Network view of results in F. J) Gene expression of selection of genes overlayed with the measured velocity vector field.

**Figure S5 - snATAC-seq cell quality control and label transfer scores. - related to Figures 5 and 6.** A) Number of fragments and Transcription Start Site (TSS) enrichment of each cell. Color represent number of cells. B) Seurat prediction score based on scRNA-seq annotation. First panel represent the maximum score for all cell types. Middle panel shows prediction score for preOPCs cluster. Last panel shows prediction score for OPCs.

**Figure S6.** PciSeq HyISS of three coronal brain slices ranging from the anterior to posterior axis of PCW 8 human forebrain, images are separated by estimated coarse cluster membership according to probabilistic mapping.

**Figure S7. PciSeq HyISS of** three **coronal brain slices ranging from the anterior to posterior axis of PCW 8 human forebrain**. A) The measured coordinates of the panel of 50 probes used for the most anterior slice. B) The measured coordinates of the panel of 50 probes used for the slice in between the most anterior and posterior. C) The measured coordinates of the panel of 50 probes used for the most posterior slice. D) DAPI staining showing the dense structure of the radial glial neuroepithelium for all three slices.

## Legends for Supplementary Tables

**Supplementary Table 1** - Differential expression test per cluster as in Seurat v4 using wilcoxon rank sum test, logFC 0.25, pct 0.25

**Supplementary Table 2** - Differential snATAC-Seq gene activity (Wilcoxon rank sum) of OPCs vs preOPCs (FDR < 0.01)

## References

Aibar, S., González-Blas, C.B., Moerman, T., Huynh-Thu, V.A., Imrichova, H., Hulselmans, G., Rambow, F., Marine, J.C., Geurts, P., Aerts, J., et al. (2017). SCENIC: Single-cell regulatory network inference and clustering. Nat. Methods 14, 1083–1086.

Arendt, D., Musser, J.M., Baker, C.V.H., Bergman, A., Cepko, C., Erwin, D.H., Pavlicev, M., Schlosser, G., Widder, S., Laubichler, M.D., et al. (2016a). The origin and evolution of cell types. Nat. Rev. Genet. 17, 744–757.

Arendt, D., Tosches, M.A., and Marlow, H. (2016b). From nerve net to nerve ring, nerve cord and brain-evolution of the nervous system. Nat. Rev. Neurosci. 17, 61–72.

Bergen, V., Lange, M., Peidli, S., Wolf, F.A., and Theis, F.J. (2019). Generalizing RNA velocity to transient cell states through dynamical modeling. 1–26.

Boulanger, J.J., and Messier, C. (2017). Doublecortin in Oligodendrocyte Precursor Cells in the Adult Mouse Brain. Front. Neurosci. 11, 143.

Brabletz, S., Bajdak, K., Meidhof, S., Burk, U., Niedermann, G., Firat, E., Wellner, U., Dimmler, A., Faller, G., Schubert, J., et al. (2011). The ZEB1/miR-200 feedback loop controls Notch signalling in cancer cells. EMBO J. 30, 770–782.

Bray, N.L., Pimentel, H., Melsted, P., and Pachter, L. (2016). Near-optimal probabilistic RNA-seq quantification. Nat. Biotechnol. 34, 525–527.

Buenrostro, J.D., Wu, B., Litzenburger, U.M., Ruff, D., Gonzales, M.L., Snyder, M.P., Chang, H.Y., and Greenleaf, W.J. (2015). Single-cell chromatin accessibility reveals principles of regulatory variation. Nature 523, 486–490.

Cao, J., Spielmann, M., Qiu, X., Huang, X., Ibrahim, D.M., Hill, A.J., Zhang, F., Mundlos, S., Christiansen, L., Steemers, F.J., et al. (2019). The single-cell transcriptional landscape of mammalian organogenesis. Nature.

Claus Stolt, C., Rehberg, S., Ader, M., Lommes, P., Riethmacher, D., Schachner, M., Bartsch, U., and Wegner, M. (2002). Terminal differentiation of myelin-forming oligodendrocytes depends on the transcription factor Sox10. Genes Dev.

Di Bella, D.J., Habibi, E., Stickels, R.R., Scalia, G., Brown, J., Yadollahpour, P., Yang, S.M., Abbate, C., Biancalani, T., Macosko, E.Z., et al. (2021). Molecular logic of cellular diversification in the mouse cerebral cortex. Nature.

Falcão, A.M., van Bruggen, D., Marques, S., Meijer, M., Jäkel, S., Agirre, E., Samudyata, Floriddia, E.M., Vanichkina, D.P., ffrench-Constant, C., et al. (2018). Disease-specific oligodendrocyte lineage cells arise in multiple sclerosis. Nat. Med. 24, 1837–1844.

Floriddia, E.M., Lourenço, T., Zhang, S., van Bruggen, D., Hilscher, M.M., Kukanja, P., Gonçalves dos Santos, J.P., Altınkök, M., Yokota, C., Llorens-Bobadilla, E., et al. (2020). Distinct oligodendrocyte populations have spatial preference and different responses to spinal cord injury. Nat. Commun. 11, 5860.

Fogarty, M., Richardson, W.D., and Kessaris, N. (2005). A subset of oligodendrocytes generated from radial glia in the dorsal spinal cord. Development 132, 1951–1959.

Genoud, S., Lappe-Siefke, C., Goebbels, S., Radtke, F., Aguet, M., Scherer, S.S., Suter, U., Nave, K.-A., and Mantei, N. (2002). Notch1 control of oligodendrocyte differentiation in the spinal cord. J. Cell Biol. 158, 709–718.

Grandbarbe, L., Bouissac, J., Rand, M., Hrabé de Angelis, M., Artavanis-Tsakonas, S., and Mohier, E. (2003). Delta-Notch signaling controls the generation of neurons/glia from neural stem cells in a stepwise process. Development 130, 1391–1402.

Gyllborg, D., Langseth, C.M., Qian, X., Choi, E., Salas, S.M., Hilscher, M.M., Lein, E.S., and Nilsson, M. (2020). Hybridization-based in situ sequencing (HybISS) for spatially resolved transcriptomics in human and mouse brain tissue. Nucleic Acids Res. 48, e112–e112.

Haghverdi, L., Büttner, M., Wolf, F.A., Buettner, F., and Theis, F.J. (2016). Diffusion pseudotime robustly reconstructs lineage branching. Nat. Methods 13, 845–848.

Haghverdi, L., Lun, A.T.L., Morgan, M.D., and Marioni, J.C. (2018). Batch effects in single-cell RNA-sequencing data are corrected by matching mutual nearest neighbors. Nat. Biotechnol. 36, 421–427.

Hammond, T.R., Gadea, A., Dupree, J., Kerninon, C., Nait-Oumesmar, B., Aguirre, A., and Gallo, V. (2014). Astrocyte-Derived Endothelin-1 Inhibits Remyelination through Notch Activation. Neuron 81, 588–602.

He, W., Ingraham, C., Rising, L., Goderie, S., and Temple, S. (2001). Multipotent Stem Cells from the Mouse Basal Forebrain Contribute GABAergic Neurons and Oligodendrocytes to the Cerebral Cortex during Embryogenesis. J. Neurosci. 21, 8854–8862.

Hochgerner, H., Lönnerberg, P., Hodge, R., Mikes, J., Heskol, A., Hubschle, H., Lin, P., Picelli, S., La Manno, G., Ratz, M., et al. (2017). STRT-seq-2i: Dual-index 5′ single cell and nucleus RNA-seq on an addressable microwell array. Sci. Rep.

Hochstim Christian, John (2009). Pax6 Controls Astrocyte Positional Identity in the Spinal Cord. California Institute of Technology.

Houart, C., Westerfield, M., and Wilson, S.W. (1998). A small population of anterior cells patterns the forebrain during zebrafish gastrulation. Nature.

Hu, Q.-D., Ang, B.-T., Karsak, M., Hu, W.-P., Cui, X.-Y., Duka, T., Takeda, Y., Chia, W., Sankar, N., Ng, Y.-K., et al. (2003). F3/Contactin Acts as a Functional Ligand for Notch during Oligodendrocyte Maturation. Cell 115, 163–175.

Huang, W., Bhaduri, A., Velmeshev, D., Wang, S., Wang, L., Rottkamp, C.A., Alvarez-Buylla, A., Rowitch, D.H., and Kriegstein, A.R. (2020). Origins and Proliferative States of Human Oligodendrocyte Precursor Cells. Cell 182, 594–608.e11.

Huynh-Thu, V.A., Irrthum, A., Wehenkel, L., and Geurts, P. (2010). Inferring Regulatory Networks from Expression Data Using Tree-Based Methods. PLoS ONE 5, e12776.

Islam, S., Zeisel, A., Joost, S., La Manno, G., Zajac, P., Kasper, M., Lönnerberg, P., and Linnarsson, S. (2014). Quantitative single-cell RNA-seq with unique molecular identifiers. Nat. Methods 11, 163–166.

Jakovcevski, I., and Zecevic, N. (2005). Sequence of oligodendrocyte development in the human fetal telencephalon. Glia 49, 480–491.

Kawaguchi, D., Furutachi, S., Kawai, H., Hozumi, K., and Gotoh, Y. (2013). Dll1 maintains quiescence of adult neural stem cells and segregates asymmetrically during mitosis. Nat. Commun. 4, 1880.

Ke, R., Mignardi, M., Pacureanu, A., Svedlund, J., Botling, J., Wählby, C., and Nilsson, M. (2013). In situ sequencing for RNA analysis in preserved tissue and cells. Nat. Methods 10, 857–860.

Kessaris, N., Fogarty, M., Iannarelli, P., Grist, M., Wegner, M., and Richardson, W.D. (2006). Competing waves of oligodendrocytes in the forebrain and postnatal elimination of an embryonic lineage. Nat. Neurosci. 9, 173–179.

Kim, H., Shin, J., Kim, S., Poling, J., Park, H.-C., and Appel, B. (2008). Notch-regulated oligodendrocyte specification from radial glia in the spinal cord of zebrafish embryos. Dev. Dyn. 237, 2081–2089.

Kirby, L., Jin, J., Cardona, J.G., Smith, M.D., Martin, K.A., Wang, J., Strasburger, H., Herbst, L., Alexis, M., Karnell, J., et al. (2019). Oligodendrocyte precursor cells present antigen and are cytotoxic targets in inflammatory demyelination. Nat. Commun. 10, 3887.

Klein, A.M., Mazutis, L., Akartuna, I., Tallapragada, N., Veres, A., Li, V., Peshkin, L., Weitz, D.A., and Kirschner, M.W. (2015). Droplet barcoding for single-cell transcriptomics applied to embryonic stem cells. Cell 161, 1187–1201.

Korsunsky, I., Millard, N., Fan, J., Slowikowski, K., Zhang, F., Wei, K., Baglaenko, Y., Brenner, M., Loh, P., and Raychaudhuri, S. (2019). Fast, sensitive and accurate integration of single-cell data with Harmony. Nat. Methods 16, 1289–1296.

Kriegstein, A., and Alvarez-Buylla, A. (2009). The Glial Nature of Embryonic and Adult Neural Stem Cells. Annu. Rev. Neurosci. 32, 149–184.

La Manno, G. (2019). From single-cell RNA-seq to transcriptional regulation. Nat. Biotechnol. 37, 1421–1422.

La Manno, G., Gyllborg, D., Codeluppi, S., Nishimura, K., Salto, C., Zeisel, A., Borm, L.E., Stott, S.R.W., Toledo, E.M., Villaescusa, J.C., et al. (2016). Molecular Diversity of Midbrain Development in Mouse, Human, and Stem Cells. Cell 167, 566–580.e19.

Lange, M., Bergen, V., Klein, M., Setty, M., Reuter, B., Bakhti, M., Lickert, H., Ansari, M., Schniering, J., Schiller, H.B., et al. (2020). CellRank for directed single-cell fate mapping (Bioinformatics).

Langseth, C.M., Gyllborg, D., Miller, J.A., Close, J.L., Long, B., Lein, E.S., Hilscher, M.M., and Nilsson, M. (2021). Comprehensive in situ mapping of human cortical transcriptomic cell types (Neuroscience).

Magri, L., Swiss, V.A., Jablonska, B., Lei, L., Pedre, X., Walsh, M., Zhang, W., Gallo, V., Canoll, P., and Casaccia, P. (2014). E2F1 coregulates cell cycle genes and chromatin components during the transition of oligodendrocyte progenitors from proliferation to differentiation. J. Neurosci. Off. J. Soc. Neurosci. 34, 1481–1493.

Marioni, J.C., and Arendt, D. (2017). How Single-Cell Genomics Is Changing Evolutionary and Developmental Biology. Annu. Rev. Cell Dev. Biol. 33, 537–553.

Marklund, U., Alekseenko, Z., Andersson, E., Falci, S., Westgren, M., Perlmann, T., Graham, A., Sundström, E., and Ericson, J. (2014). Detailed Expression Analysis of Regulatory Genes in the Early Developing Human Neural Tube. Stem Cells Dev. 23, 5–15.

Marques, S., van Bruggen, D., Vanichkina, D.P., Floriddia, E.M., Munguba, H., Väremo, L., Giacomello, S., Falcão, A.M., Meijer, M., Björklund, Å.K., et al. (2018). Transcriptional Convergence of Oligodendrocyte Lineage Progenitors during Development. Dev. Cell 46, 504–517.e7.

McClain, C.R., Sim, F.J., and Goldman, S.A. (2012). Pleiotrophin Suppression of Receptor Protein Tyrosine Phosphatase- / Maintains the Self-Renewal Competence of Fetal Human Oligodendrocyte Progenitor Cells. J. Neurosci. 32, 15066–15075.

Naka, H., Nakamura, S., Shimazaki, T., and Okano, H. (2008). Requirement for COUP-TFI and II in the temporal specification of neural stem cells in CNS development. Nat. Neurosci. 11, 1014–1023.

Nowakowski, T.J., Bhaduri, A., Pollen, A.A., Alvarado, B., Mostajo-Radji, M.A., Di Lullo, E., Haeussler, M., Sandoval-Espinosa, C., Liu, S.J., Velmeshev, D., et al. (2017). Spatiotemporal gene expression trajectories reveal developmental hierarchies of the human cortex. Science 358, 1318–1323.

Park, H.-C. (2005). Oligodendrocyte Specification in Zebrafish Requires Notch-Regulated Cyclin-Dependent Kinase Inhibitor Function. J. Neurosci. 25, 6836–6844.

Polioudakis, D., Torre-ubieta, L.D., Langerman, J., Gerstein, M.B., Plath, K., Geschwind, D.H., Polioudakis, D., Torre-ubieta, L.D., Langerman, J., Elkins, A.G., et al. (2019). NeuroResource A Single-Cell Transcriptomic Atlas of Human Neocortical Development during Mid-gestation NeuroResource A Single-Cell Transcriptomic Atlas of Human Neocortical Development during Mid-gestation. Neuron 1–17.

Pollen, A.A., Nowakowski, T.J., Chen, J., Retallack, H., Sandoval-Espinosa, C., Nicholas, C.R., Shuga, J., Liu, S.J., Oldham, M.C., Diaz, A., et al. (2015). Molecular Identity of Human Outer Radial Glia during Cortical Development. Cell 163, 55–67.

Potzner, M.R., Griffel, C., Lütjen-Drecoll, E., Bösl, M.R., Wegner, M., and Sock, E. (2007). Prolonged Sox4 Expression in Oligodendrocytes Interferes with Normal Myelination in the Central Nervous System. Mol. Cell. Biol. 27, 5316–5326.

Qian, X., Harris, K.D., Hauling, T., Nicoloutsopoulos, D., Muñoz-Manchado, A.B., Skene, N., Hjerling-Leffler, J., and Nilsson, M. (2020). Probabilistic cell typing enables fine mapping of closely related cell types in situ. Nat. Methods 17, 101–106.

R Development Core Team (2010). a language and environment for statistical computing: reference index (Vienna: R Foundation for Statistical Computing).

Rayon, T., Maizels, R.J., Barrington, C., and Briscoe, J. (2021). Single cell transcriptome profiling of the human developing spinal cord reveals a conserved genetic programme with human specific features (Developmental Biology).

Rosenbaum, J.N., Guo, Z., Baus, R.M., Werner, H., Rehrauer, W.M., and Lloyd, R.V. (2015). INSM1: A Novel Immunohistochemical and Molecular Marker for Neuroendocrine and Neuroepithelial Neoplasms. Am. J. Clin. Pathol. 144, 579–591.

Rowitch, D.H., and Kriegstein, A.R. (2010). Developmental genetics of vertebrate glial-cell specification. Nature 468, 214–222.

Sim, F.J., McClain, C.R., Schanz, S.J., Protack, T.L., Windrem, M.S., and Goldman, S.A. (2011). CD140a identifies a population of highly myelinogenic, migration-competent and efficiently engrafting human oligodendrocyte progenitor cells. Nat. Biotechnol. 29, 934–941.

Speir, M.L., Bhaduri, A., Markov, N.S., Moreno, P., Nowakowski, T.J., Papatheodorou, I., Pollen, A.A., Raney, B.J., Seninge, L., Kent, W.J., et al. (2021). UCSC Cell Browser: visualize your single-cell data. Bioinformatics btab503.

Stolt, C.C., and Wegner, M. (2010). SoxE function in vertebrate nervous system development.

Stuart, T., Butler, A., Hoffman, P., Hafemeister, C., Papalexi, E., Mauck, W.M., Hao, Y., Stoeckius, M., Smibert, P., and Satija, R. (2019). Comprehensive Integration of Single-Cell Data. Cell 177, 1888–1902.e21.

Tavano, S., Taverna, E., Kalebic, N., Haffner, C., Namba, T., Dahl, A., Wilsch-Bräuninger, M., Paridaen, J.T.M.L., and Huttner, W.B. (2018). Insm1 Induces Neural Progenitor Delamination in Developing Neocortex via Downregulation of the Adherens Junction Belt-Specific Protein Plekha7. Neuron 97, 1299–1314.e8.

de la Torre-Ubieta, L., Stein, J.L., Won, H., Opland, C.K., Liang, D., Lu, D., and Geschwind, D.H. (2018). The Dynamic Landscape of Open Chromatin during Human Cortical Neurogenesis. Cell 172, 289–304.e18.

Ugrinova, I., Pashev, I.G., and Pasheva, E.A. (2009). Nucleosome binding properties and Co-remodeling activities of native and in vivo acetylated HMGB-1 and HMGB-2 proteins. Biochemistry 48, 6502–6507.

Van de Sande, B., Flerin, C., Davie, K., De Waegeneer, M., Hulselmans, G., Aibar, S., Seurinck, R., Saelens, W., Cannoodt, R., Rouchon, Q., et al. (2020). A scalable SCENIC workflow for single-cell gene regulatory network analysis. Nat. Protoc. 15, 2247–2276.

Wang, C., Zhang, C.-J., Martin, B.N., Bulek, K., Kang, Z., Zhao, J., Bian, G., Carman, J.A., Gao, J., Dongre, A., et al. (2017). IL-17 induced NOTCH1 activation in oligodendrocyte progenitor cells enhances proliferation and inflammatory gene expression. Nat. Commun. 8, 15508.

Wang, S., Sdrulla, A.D., diSibio, G., Bush, G., Nofziger, D., Hicks, C., Weinmaster, G., and Barres, B.A. (1998). Notch Receptor Activation Inhibits Oligodendrocyte Differentiation. Neuron 21, 63–75.

Windrem, M.S., Nunes, M.C., Rashbaum, W.K., Schwartz, T.H., Goodman, R.A., McKhann, G., Roy, N.S., and Goldman, S.A. (2004). Fetal and adult human oligodendrocyte progenitor cell isolates myelinate the congenitally dysmyelinated brain. Nat. Med. 10, 93–97.

Windrem, M.S., Schanz, S.J., Zou, L., Chandler-Militello, D., Kuypers, N.J., Nedergaard, M., Lu, Y., Mariani, J.N., and Goldman, S.A. (2020). Human Glial Progenitor Cells Effectively Remyelinate the Demyelinated Adult Brain. Cell Rep. 31, 107658.

Winkler, C.C., Yabut, O.R., Fregoso, S.P., Gomez, H.G., Dwyer, B.E., Pleasure, S.J., and Franco, S.J. (2018). The Dorsal Wave of Neocortical Oligodendrogenesis Begins Embryonically and Requires Multiple Sources of Sonic Hedgehog. J. Neurosci. 38, 5237–5250.

Wittstatt, J., Reiprich, S., and Küspert, M. (2019). Crazy Little Thing Called Sox—New Insights in Oligodendroglial Sox Protein Function. Int. J. Mol. Sci. 20, 2713.

Won, H., de la Torre-Ubieta, L., Stein, J.L., Parikshak, N.N., Huang, J., Opland, C.K., Gandal, M.J., Sutton, G.J., Hormozdiari, F., Lu, D., et al. (2016). Chromosome conformation elucidates regulatory relationships in developing human brain. Nature 538, 523–527.

Wood, S.N., Pya, N., and Säfken, B. (2016). Smoothing Parameter and Model Selection for General Smooth Models. J. Am. Stat. Assoc. 111, 1548–1563.

Zeisel, A., Hochgerner, H., Lönnerberg, P., Johnsson, A., Memic, F., van der Zwan, J., Häring, M., Braun, E., Borm, L.E., La Manno, G., et al. (2018). Molecular Architecture of the Mouse Nervous System. Cell 174, 999–1014.e22.

Zhang, Y., Hoxha, E., Zhao, T., Zhou, X., and Alvarez-Bolado, G. (2017). Foxb1 Regulates Negatively the Proliferation of Oligodendrocyte Progenitors. Front. Neuroanat. 11, 53.

Zhu, Y., Sousa, A.M.M., Gao, T., Skarica, M., Li, M., Santpere, G., Esteller-Cucala, P., Juan, D., Ferrández-Peral, L., Gulden, F.O., et al. (2018). Spatiotemporal transcriptomic divergence across human and macaque brain development. Science 362, eaat8077.

Single-cell reconstruction of developmental trajectories during zebrafish embryogenesis.

