## Supplementary Figure S1-S7 for "Developmental landscape of human forebrain at a single-cell level unveils early waves of oligodendrogenesis"

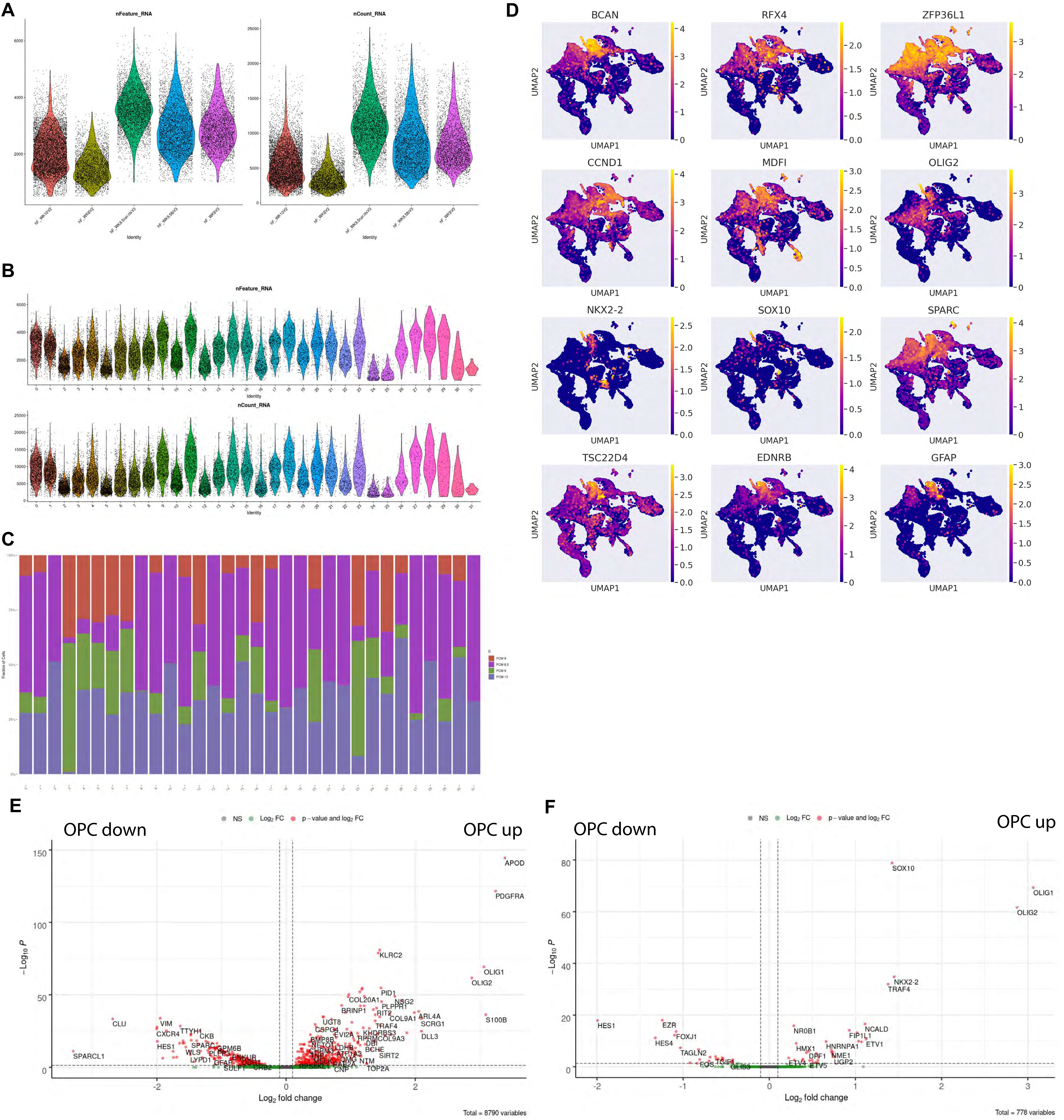


Figure S1. A) Violin plot showing the distribution of the number of counts and number of features over all samples. B) Violin plot showing the distribution of the number of counts and number of features overall clusters. C) Barplot showing the distribution of age over the clusters. D) UMAP plots of selected genes. E) Volcano-plot showing differential expression results for preOPC compared to OPC. F) Volcano-plot showing differential expression results for preOPC compared to OPC but only annotated transcriptionfactors and co-factors.

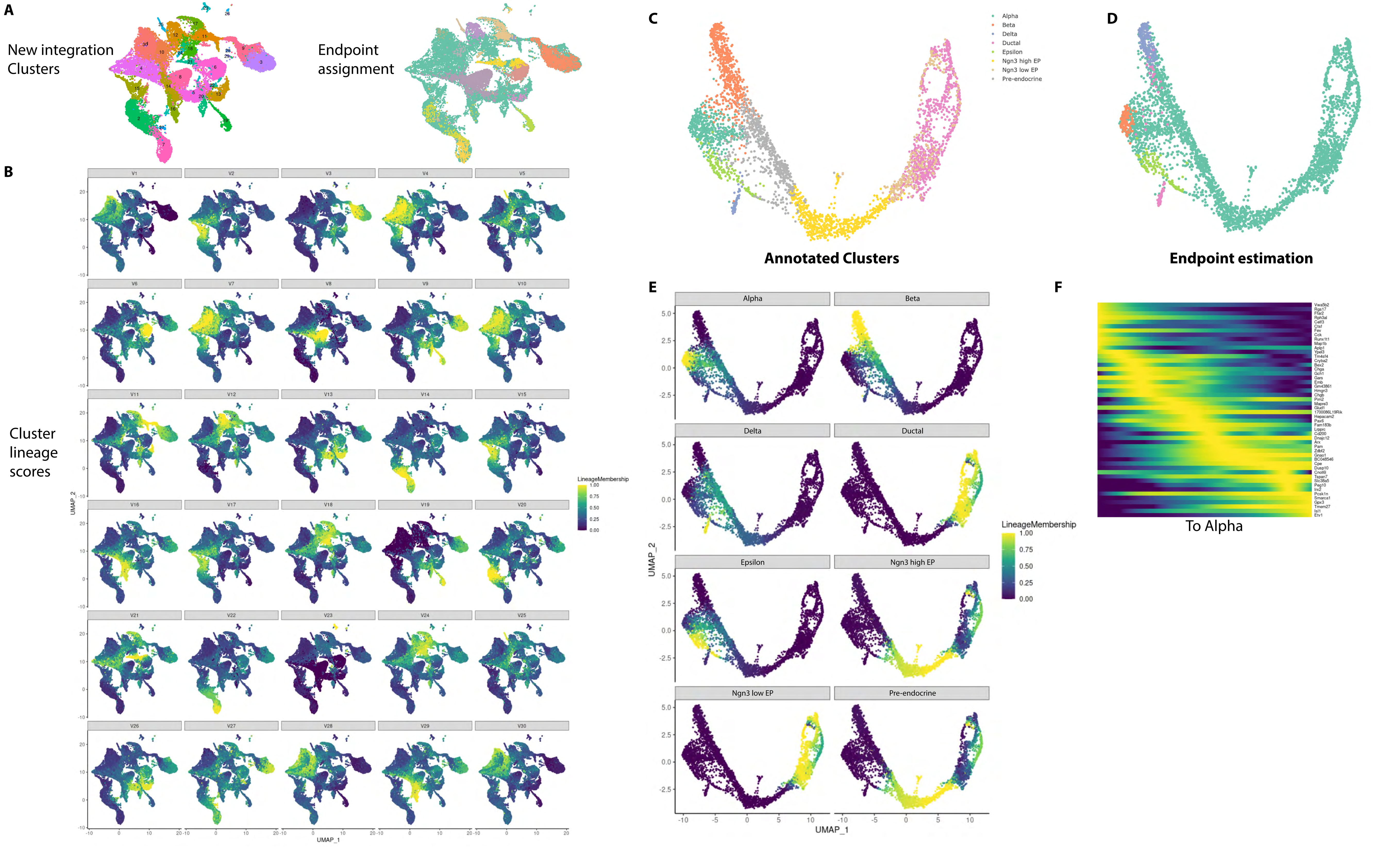


Figure S2. A) Left, UMAP showing the clustering performed for the lineage analysis after integration. Right, Endpoint assignment results of the lineage analysis. B) Lineage membership scores calculated for each cluster (softscaled (exp)0.1). C) UMAP of the benchmark test set as from the pancreatic dataset. D) Endpoint assignment results of the lineage analysis on the pancreatic dataset. E) Lineage membership scores for all clusters of the pancreatic dataset. F) Gene ordering and cell ordering towards the Alpha lineage in the pancreatic dataset.

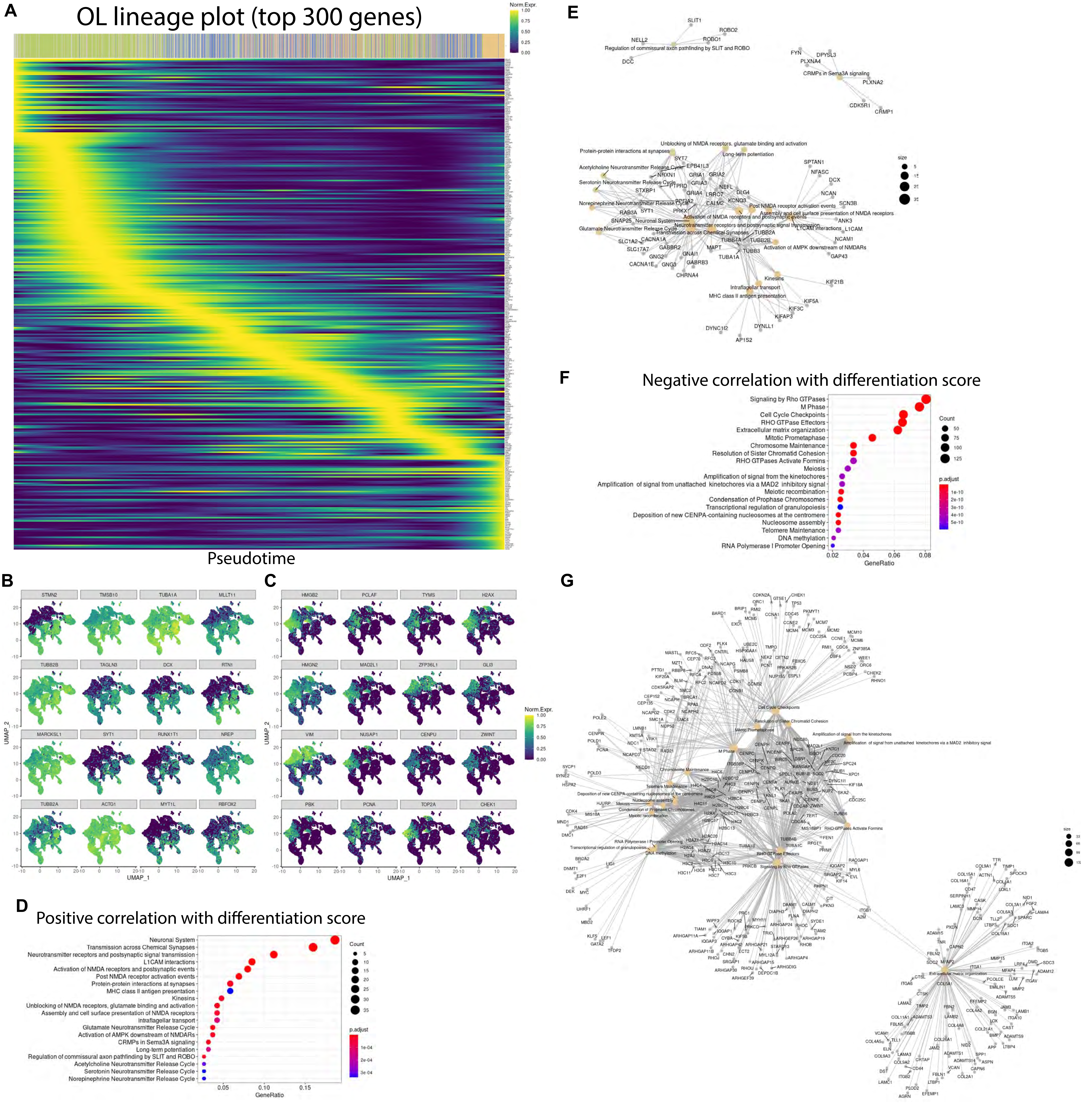


Figure S3. A) Gene ordering and cell ordering towards the Oligodendrocyte lineage. B) Top 16 genes positively correlating with the differentiation score over the whole dataset (Pearson R, Fishertest, FDR 5%). C) Top 16 genes negatively correlating with the differentiation score over the whole dataset (Pearson R, Fishertest, FDR 5%). D) Reactome pathway analysis over the significantly positively correlating genes over the whole dataset (Pearson R, Fishertest, FDR 5%). E) Network view of results in D. F) Reactome pathway analysis over the significantly negatively correlating genes over the whole dataset (Pearson R, Fishertest, FDR 5%). G) Network view of results in F.

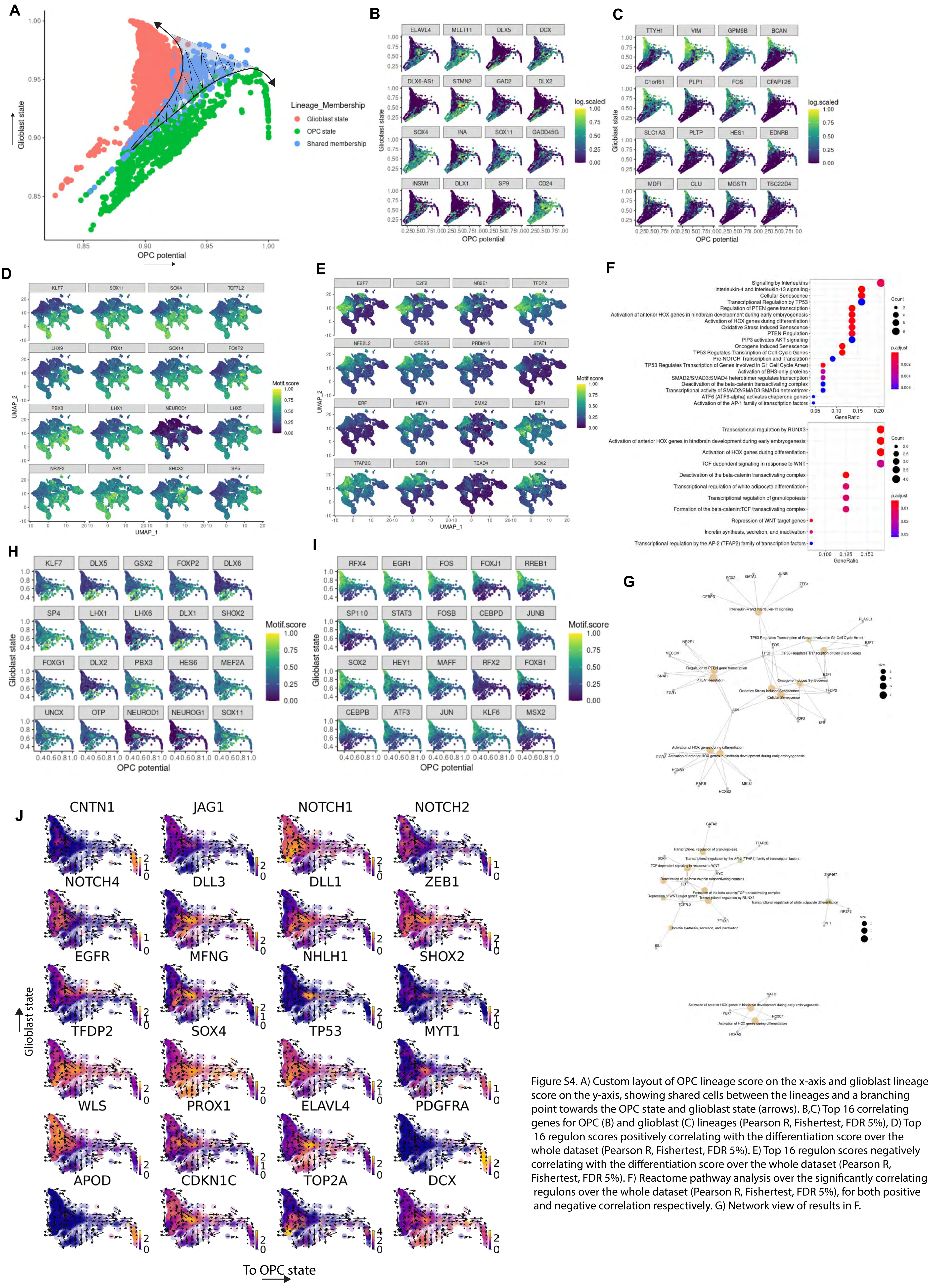

Figure S4. A) Custom layout of OPC lineage score on the x-axis and glioblast lineage score on the y-axis, showing shared cells between the lineages and a branching point towards the OPC state and glioblast state (arrows). B,C) Top 16 correlating genes for OPC (B) and glioblast (C) lineages (Pearson R, Fishertest, FDR 5%), D) Top 16 regulon scores positively correlating with the differentiation score over the whole dataset (Pearson R, Fishertest, FDR 5%). E) Top 16 regulon scores negatively correlating with the differentiation score over the whole dataset (Pearson R, Fishertest, FDR 5%). F) Reactome pathway analysis over the significantly correlating regulons over the whole dataset (Pearson R, Fishertest, FDR 5%), for both positive and negative correlation respectively. G) Network view of results in F.

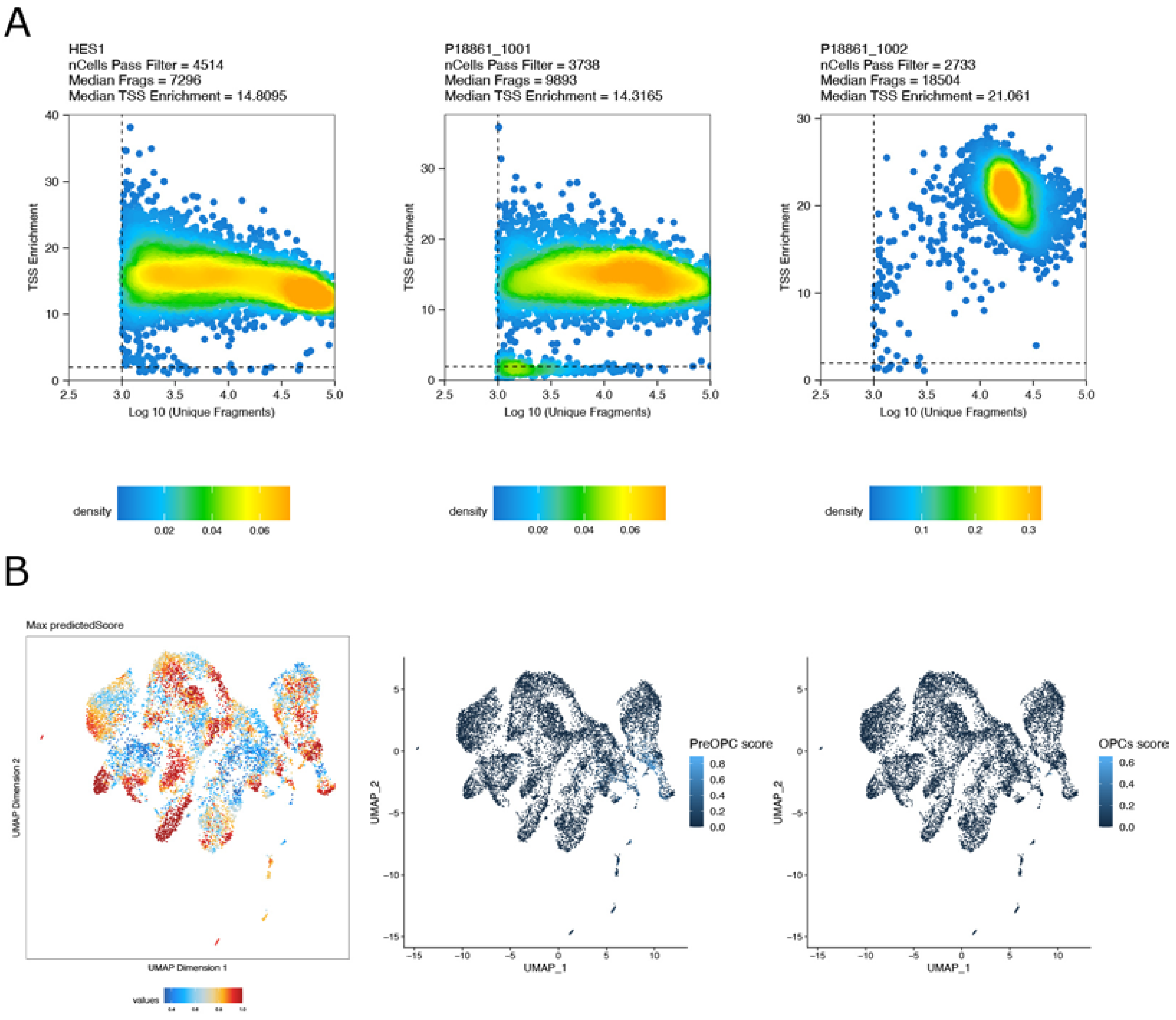


Figure 5S - snATAC-seq cell quality control and label transfer scores. A) Number of fragments and Transcription Start Site (TSS) enrichment of each cell. Color represent number of cells. B) Seurat prediction score based on scRNA-seq annotation. First panel represent the maximum score for all cell types. Middle panel shows prediction score for preOPCs cluster. Last panel shows prediction score for OPCs.

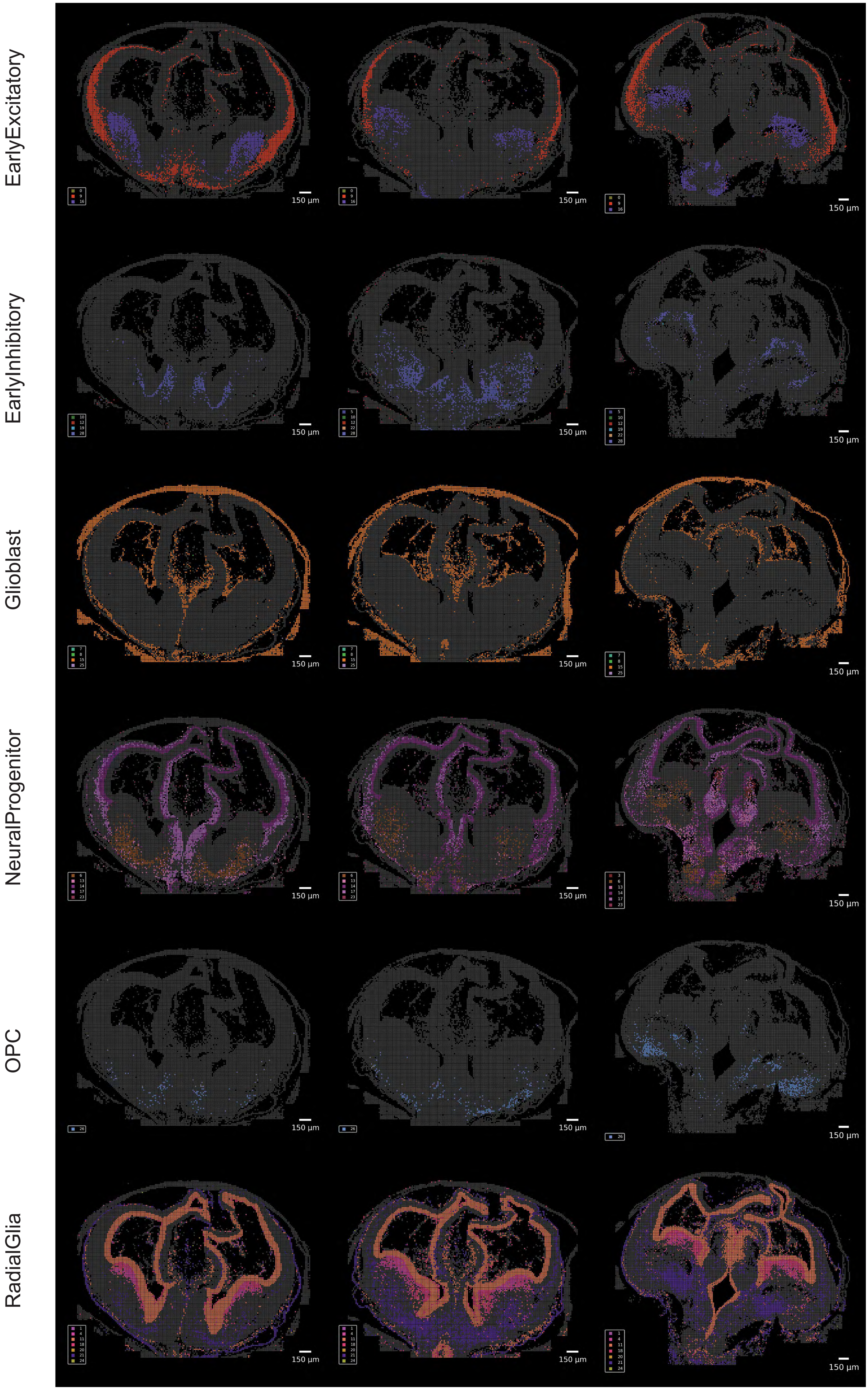


Figure S6. PciSeq HyISS of three coronal brain slices ranging from the anterior to posterior axis of PCW 8 human forebrain, images are separated by estimated coarse cluster membership according to probabilistic mapping.

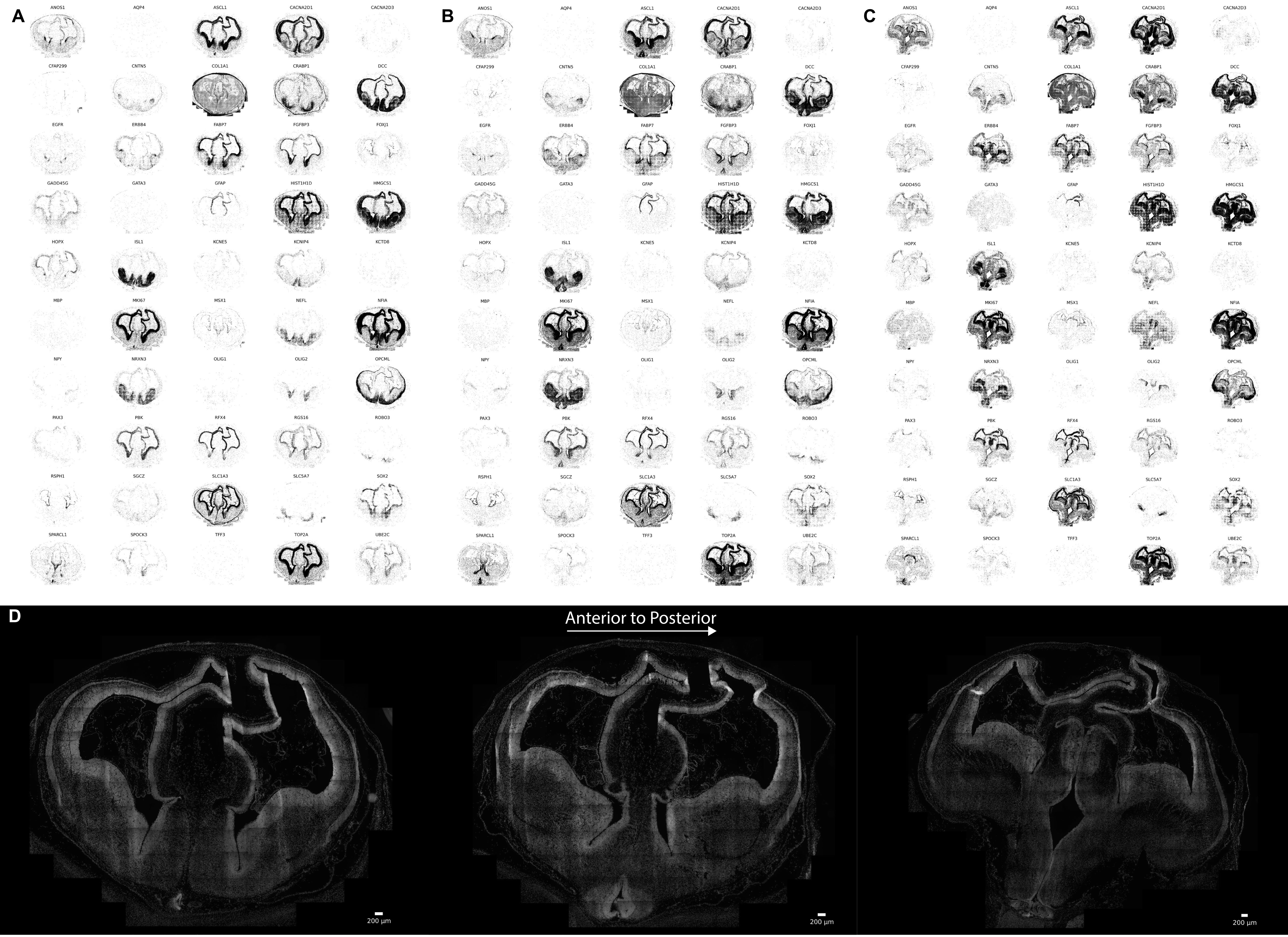


Figure S7. PciSeq HyISS of three coronal brain slices ranging from the anterior to posterior axis of PCW 8 human forebrain. A) The measured coordinates of the panel of 50 probes used for the most anterior slice. B) The measured coordinates of the panel of 50 probes used for the slice in between the most anterior and posterior. C) The measured coordinates of the panel of 50 probes used for the most posterior slice. D) DAPI staining showing the dense structure of the radial glial neuroepithelium for all three slices.
